# Rapid direct neuronal reprogramming of human dental pulp stem cells

**DOI:** 10.64898/2026.07.24.740560

**Authors:** Anna A. Abbas, Chandramouli Muralidharan, Bendegúz Sramkó, Melinda E. Gazdik, Enikő Zakar-Polyák, Brigitta Varga, Kinga Vörös, Balázs Kis, Ágnes Varga, Jenny G. Johansson, Kristóf Kádár, Zsuzsanna Darula, Csaba Kerepesi, Karri Lamsa, Attila Szűcs, Gábor Varga, Ákos Zsembery, Anna Földes, Karolina Pircs

**Affiliations:** Institute of Clinical Pathophysiology, Semmelweis University, 1094 Budapest, Hungary; Hungarian Centre of Excellence for Molecular Medicine - Semmelweis University (HCEMM-SU), Neurobiology and Neurodegenerative Diseases Research Group, 1094 Budapest, Hungary; Laboratory of Molecular Neurogenetics, Department of Experimental Medical Science, Wallenberg Neuroscience Center and Lund Stem Cell Center, BMC A11, Lund University, 22184 Lund, Sweden; HUN-REN-SZTAKI-SU Rejuvenation Research Group, HUN-REN Office for Supported Research Groups (TKI), 1052 Budapest, Hungary; Department of Physiology and Neurobiology, Institute of Biology, Eötvös Loránd University, 1117 Budapest, Hungary; Institute for Computer Science and Control (SZTAKI), Hungarian Research Network (HUN-REN), 1111 Budapest, Hungary; Doctoral School of Informatics, Faculty of Informatics, Eötvös Loránd University, Budapest 1053, Hungary; Department of Oral Biology, Faculty of Dentistry, Semmelweis University, 1089 Budapest, Hungary; Single Cell Omics Advanced Core Facility, Hungarian Centre of Excellence for Molecular Medicine, 6728 Szeged, Hungary; Laboratory of Proteomics, Complex Molecular and Cell Biology Service Centre, HUN-REN Biological Research Centre, 6726 Szeged, Hungary; Hungarian Center of Excellence for Molecular Medicine Research Group for Human neuron physiology and therapy, 6728 Szeged, Hungary; Department of Physiology, Anatomy and Neuroscience, University of Szeged, 6726 Szeged, Hungary; Centre for Translational Medicine, Semmelweis University, 1085 Budapest, Hungary

**Keywords:** induced neurons, direct neuronal reprogramming, dental pulp stem cells, single-nucleus RNA sequencing, neuronal heterogeneity

## Abstract

Direct neuronal reprogramming offers an alternative to induced pluripotent stem cell-based differentiation by converting somatic cells directly into neurons without passage through pluripotency. However, commonly used fibroblast-based protocols are often slow and inefficient. Here, we evaluated human dental pulp stem cells (DPSCs), which originate from the cranial neural crest and possess intrinsic neurogenic potential, as a developmentally relevant source of induced neurons (iNs). Using an all-in-one lentiviral vector, we converted DPSCs into iNs within 17 days, compared with 28 days for fibroblasts reprogrammed with the same vector, and achieved significantly higher neuronal purity under the respective established protocols. Multi-omic profiling revealed coordinated suppression of mesenchymal and cell-cycle programs and induction of neuronal, synaptic, and metabolic pathways. Single-nucleus RNA sequencing resolved fibroblast-like, transitional, maturing neuronal, GABAergic-like, and alternative fates, while trajectory inference suggested divergent neuronal and non-neuronal conversion paths. Whole-cell recordings showed that a subset of DPSC-iNs developed early neuronal excitability and voltage-gated inward and outward currents. Together, our findings establish DPSCs as an accessible and developmentally relevant source for rapid direct neuronal conversion. This integrated molecular, single-nucleus, and electrophysiological characterization defines the cellular heterogeneity of DPSC-to-neuron reprogramming and provides a framework for protocol refinement and future patient-specific disease modeling.

## INTRODUCTION

Direct reprogramming, also referred to as transdifferentiation, is the process of converting one somatic cell type into another through the forced expression of lineage-specific transcription factors, microRNAs, or small molecules that reshape the cell’s transcriptional, epigenetic, and functional identity^1–3^. Unlike reprogramming via induced pluripotency, direct conversion bypasses intermediate progenitor or pluripotent states, highlighting the influence of a cell’s developmental origin on both purity and outcome. This approach has enabled the generation of *in vitro* neurons that retain epigenetic alterations acquired during pathological processes or normal aging^4,5^.

Compared with induced pluripotent stem cell (iPSC)-derived neurons, direct neuronal reprogramming retains age-associated epigenetic and transcriptomic signatures, including DNA methylation patterns. This is particularly relevant for modeling human aging and late-onset neurodegenerative diseases, where donor age is a critical factor^6–8^. The development of fibroblast-to-neuron (iN) protocols was pivotal in this field, and dermal fibroblasts remain the most widely used donor cell type. However, their strong mesenchymal identity restricts neuronal lineage trajectories, limits neuronal subtype diversity, and contributes to relatively low efficiency and incomplete maturation^9–12^. These limitations have prompted exploration of alternative donor cell sources with developmental features more permissive to neuronal fate^13–15^.

Human dental pulp stem cells (DPSCs) represent a promising alternative. Unlike fibroblasts, DPSCs originate from the cranial neural crest - a lineage with established developmental relevance to the nervous system and well-documented cellular plasticity^16–19^. Tooth development arises from reciprocal interactions between neural crest-derived mesenchyme and oral epithelium, giving rise to a heterogeneous pulp tissue containing mesenchymal, epithelial, endothelial, and fibroblast-like cells. From this niche, DPSCs can be readily isolated, typically from exfoliated deciduous teeth or extracted third molars (wisdom teeth). The pulp stroma harbors mesenchymal stem cells (MSCs) with distinct phenotypic and functional features, contributing to the regenerative potential of the tissue^20,21^. Importantly, over the past two decades, several studies have shown that DPSCs can adopt neural characteristics *in vitro* when exposed to relevant growth factors or small molecules^22–24^. This intrinsic neurogenic potential likely reflects their cranial neural crest origin and epigenetic plasticity. Recent studies have used single-cell transcriptomics to resolve neural-induction states in DPSC cultures and have demonstrated electrophysiologically active neuron-like derivatives generated from DPSCs^25,26^. The present study extends these observations by combining transcription factor-mediated direct conversion with integrated bulk and single-nucleus transcriptomics, proteomics, DNA methylation profiling, and electrophysiology^25,26^. Nevertheless, like other MSCs, DPSCs are heterogeneous in culture, with variability in proliferation, differentiation potential, and marker expression. The absence of reliable DPSC surface markers for isolating functionally defined subpopulations remains a barrier to both mechanistic studies and clinical translation^27–29^.

Here, we show that human DPSCs can be directly converted into functional iNs with markedly accelerated kinetics compared to fibroblast-derived iNs (17 vs. 28 days). Using multi-omics - including transcriptomics (bulk RNAseq), proteomics, DNA methylation profiling, and single-nucleus RNA sequencing (snRNAseq) - we characterized the fate of reprogrammed DPSCs. The results consistently revealed a post-mitotic neuronal profile, while trajectory inference suggested a putative transition toward a GABAergic-like neuronal state. In addition, we demonstrate early functional maturation of DPSC-iNs using whole-cell current- and voltage-clamp electrophysiological recordings. Together, this work establishes DPSCs as a developmentally primed, readily accessible donor source for direct neuronal reprogramming, and provides the first integrated characterization of their lineage potential, opening new avenues for patient-specific modeling of human neurodevelopment and age-related neurodegenerative disease.

## MATERIALS AND METHODS

### DPSC cultures

In this study we used DPSCs from four surgically removed wisdom teeth from four different donors. DPSCs were obtained via surgical wisdom tooth removal, as previously described^23^. The wisdom teeth were obtained at the Institute of Oral Biology, Semmelweis University and used under local ethical approvals (25459-4/2019/EKU). The cohort included both male and female donors aged 18-29 years, and the cells were used between passages 2 and 4 (Table 1).

**Table 1.** Summary of DPSC donors. Age (years), Sex (F = female; M = male).

| Donor ID | Age | Sex | Passage number |
| --- | --- | --- | --- |
| DPSC1 | 18 | F | 2 |
| DPSC2 | 24 | M | 3 |
| DPSC3 | 28 | F | 3 |
| DPSC4 | 29 | F | 4 |

### Fibroblast lines

In this study, adult human dermal fibroblasts (FB) from four independent donors were used. The fibroblast lines were obtained from the John van Geest Centre for Brain Repair, University of Cambridge, UK, and from the HCEMM-USZ Skin Research Group, Hungary. The four donors, comprising both male and female individuals, ranged in age from 27 to 31 years (Table 2).

**Table 2.** Summary of FB donors. Age (years), Sex (F = female; M = male).

| Donor ID | Age | Sex | Passage number |
| --- | --- | --- | --- |
| FB1 | 27 | M | P10 |
| FB2 | 27 | M | P9 |
| FB3 | 29 | F | P11 |
| FB4 | 31 | F | P12 |

### Cell culture

Adult DPSCs were cultured in MEM alpha modified medium (Capricorn), GlutaMax Supplement (Gibco), enriched with 10% foetal bovine serum (FBS, Cytiva HyClone) and 1% Penicillin-Streptomycin (Gibco). Following an established protocol, cells were passaged upon reaching 80 - 90% confluency following the protocol described previously and used for further experiments^30^. Adult human dermal fibroblast lines were maintained in high-glucose DMEM containing GlutaMAX Supplement (Gibco), supplemented with 10% fetal bovine serum (Cytiva HyClone) and 1% penicillin/streptomycin (Gibco) and coated with 0.1% gelatin (Sigma). Cells were passaged at 80–90% confluency and routinely monitored for mycoplasma contamination based on the Uphoff and Dexter PCR method.

### Lentiviral production

For neuronal conversion we used the all-in-one transfer vector LV.U6.shREST1.U6.shREST2.hPGK.BRN2.hPGK.ASCL1.WPRE (Addgene plasmid #101852)^31^. Third generation lentiviral vectors were produced in HEK293T cells, and the virus titer was determined with RT-qPCR as previously described^32^. For DPSC neuron conversion, two viral batches were used at an MOI of 20, with titers ranging from 3.33 × 10^8^ and 1.44 × 10^9^. For fibroblast conversion, the lentiviral vector was applied at an MOI of 10 using a virus titer of 3.58 × 10^8^.

### Neuronal conversion from DPSC

24-well plates (Nunc Cell-Culture Treated Multidishes), T25 or T75 flasks (Sarstedt) were coated with 0.1% gelatin (Sigma) to enhance cell adhesion. DPSCs were seeded when 80-90% confluency was achieved, corresponding to a density of 20,000 - 35,000 cells/well (10,000-15,000 cells/cm²) for 24-well plates and 100,000-150,000 cells/T25 flask (4,000-6,000 cells/cm²) and for T75 300,000-350,000 cells/flask (4,000-4,700 cells/cm²). During expansion cells were cultured in MEM alpha modified medium (Capricorn) (Table 3). The following day (Day 1), cells were transduced with the all-in-one neuronal lentiviral vector at MOI 20. Transduction steps were conducted in a Neurobasal Plus medium (Thermo Fisher). Throughout the conversion period, cells were maintained in Neuronal conversion medium (Table 3). The conversion was performed as previously described until day 17 when the cells were either fixed for ICC staining or harvested for downstream analyses^33^.

**Table 3.** Composition of cellular media. Summary of stock concentrations and working concentrations of the components applied for DPSC medium, freezing medium, transduction medium, and neuronal conversion medium.

| ***DPSC medium*** |  |  |
| --- | --- | --- |
| **Component** | **Stock cc** | **Working cc** |
| MEM alpha modified medium | N/A | N/A |
| GlutaMax Supplement | N/A | 1% |
| Penicillin/Streptomycin | 10,000 U/mL | 100 µg/mL |
| FBS | N/A | 10% |
| ***Freezing medium*** |  |  |
| **Component** | **Stock cc** | **Working cc** |
| MEM alpha modified medium | N/A | 45% |
| FBS | N/A | 45% |
| DMSO | N/A | 10% |
| ***Transduction medium*** |  |  |
| **Component** | **Stock cc** | **Working cc** |
| MEM alpha modified medium | N/A | N/A |
| GlutaMax Supplement | N/A | 1% |
| Penicillin/Streptomycin | 10,000 U/mL | 100 µg/mL |
| FBS | N/A | 10% |
| ***Neuronal conversion medium*** |  |  |
| **Component** | **Stock cc** | **Working cc** |
| Neurobasal Plus medium | N/A | N/A |
| Penicillin/Streptomycin | 10,000 U/mL | 100 µg/mL |
| CHIR99021 | 10 mM | 2 µM |
| SB-431542 | 20 mM | 10 µM |
| Noggin | 100 µg/mL | 0.5 µg/mL |
| LDN-193189 | 10 mM | 0.5 µM |
| VPA | 1 M | 1 mM |
| LM-22A4 | 20 mM | 2 µM |
| GDNF | 20 µg/mL | 2 ng/mL |
| NT3 | 10 µg/mL | 10 ng/mL |
| db-cAMP | 50 mM | 0.5 mM |

### Neuronal conversion from FB

Fibroblasts were seeded at 80–90% confluency into 0.1% gelatin-coated (Sigma), 96-well plates (Thermo Fisher Scientific) at a density of 8,000 cells per well (25,000 cells/cm²). After overnight attachment, cells were transduced in fibroblast medium with the same all-in-one neuronal lentiviral vector used for DPSC conversion at an MOI of 10. Neuronal conversion was carried out according to the published protocols^32,31^. Cells were maintained until day 27–28 and subsequently fixed for immunocytochemical evaluation.

### Brightfield Microscopy

Bright-field images in Fig. 1b acquired using a Zeiss Axiovert 40 CFL inverted microscope at 10× and 20× magnification to document cellular conversion.

**Figure 1.**
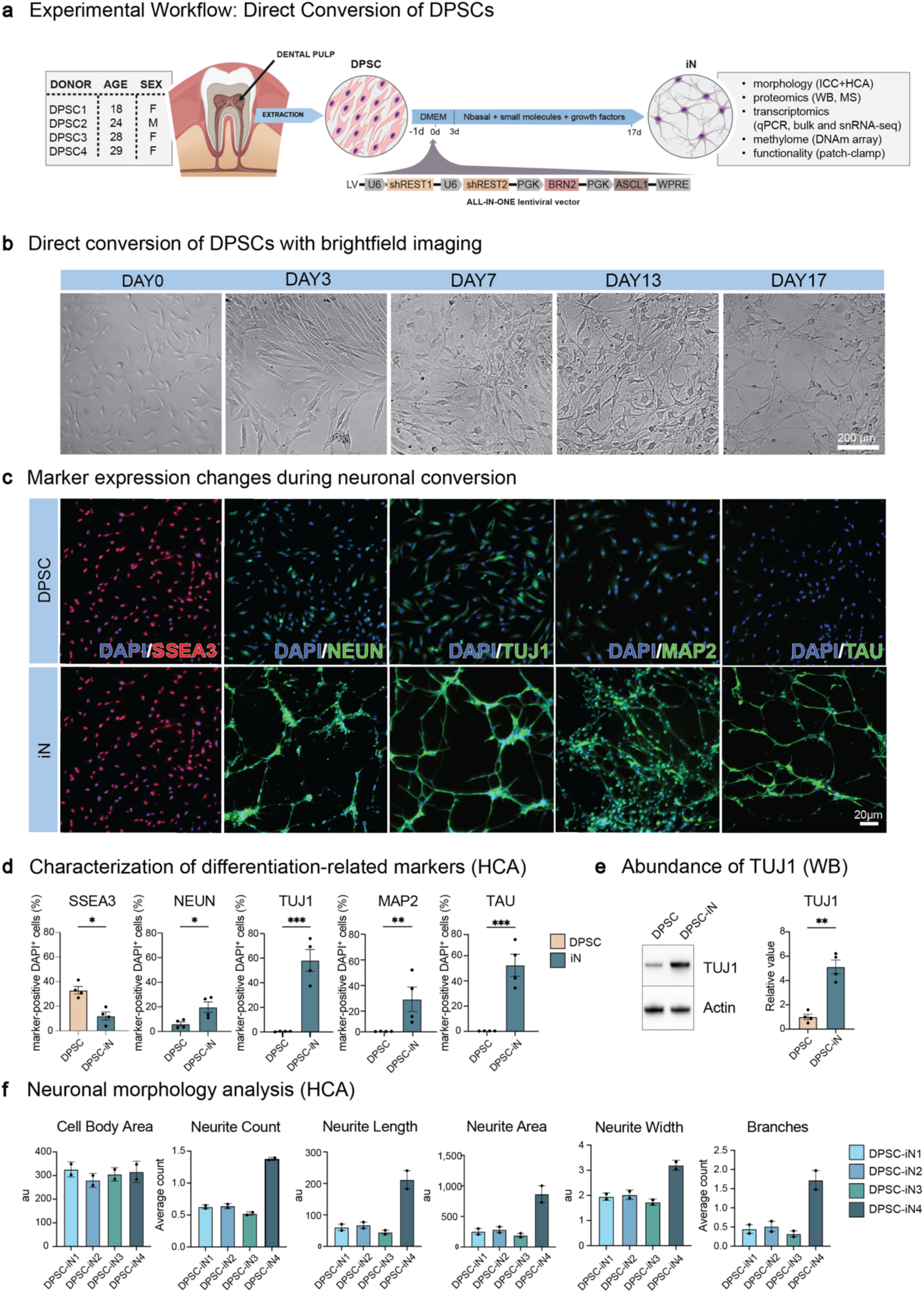
Human DPSCs readily convert into DPSC-iNs with high purity. **a:** Overview of DPSC-to-iN conversion workflow using an all-in-one lentiviral vector. **b:** Brightfield images showing morphological progression from spindle-shaped DPSCs (day 0) to neuron-like cells with neurites (day 7 - 17). Scale bar: 200 μm. **c:** Representative ICC images of DPSCs and DPSC-iNs stained for SSEA3, NeuN, TUJ1, MAP2 and TAU. **d:** HCA quantification of marker expression (% of DAPI⁺ cells). DPSC-iNs show significantly reduced SSEA3 and significantly increased NeuN, TUJ1, MAP2 and TAU (n = 4 donors). **e:** TUJ1 protein levels by WB show significantly higher TUJ1 abundance in DPSC-iNs compared to DPSCs (n = 4 donors). **f:** HCA-based morphological profiling of TAU⁺ neurons (n = 4 donors, 16 wells analyzed in total for SSEA3, NeuN, TUJ1, MAP2 and TAU). Scale bar: 20 μm. All results are shown as mean ± SEM. A one-tailed ratio paired t-test was used for panels d and e. Panel f shows descriptive donor-level morphological measurements. Significance: *p < 0.05, **p < 0.01, ***p < 0.001. Abbreviations: DPSC, dental pulp stem cell; FB, fibroblast; iN, induced neuron; sh, short hairpin; REST1/2, RE1-silencing transcription factor 1/2; PGK, phosphoglycerate kinase promoter; BRN2, POU3F2; ASCL1, achaete-scute homolog 1; WPRE, woodchuck hepatitis virus post-transcriptional regulatory element; SSEA3, stage-specific embryonic antigen 3; MAP2, microtubule-associated protein 2; NeuN/RBFOX3, RNA-binding protein fox-1 homolog 3; DMEM, Dulbecco’s modified Eagle medium.

### Immunocytochemistry (ICC)

Native and converted cells were processed for ICC following a modified protocol from our earlier work^31^. Cells were fixed in 4% paraformaldehyde (PFA) for 10 min at room temperature and rinsed twice with Dulbecco’s phosphate-buffered saline (DPBS, Capricorn Scientific). Permeabilization was achieved with 0.1% Triton X-100 in 0.1 M PBS for 10 min. Non-specific binding was blocked by incubating samples in 5% donkey serum prepared in 0.1 M PBS for at least 30 min. Primary antibodies, diluted in blocking solution, were applied overnight at 4 °C (Table 4). The following day, cells were washed twice with DPBS and incubated with fluorophore-conjugated secondary antibodies in blocking buffer for 2 h at room temperature (Table 4). Nuclei were counterstained with DAPI (1:1,000, 15 min), and samples were washed with DPBS before imaging. Fluorescence images were acquired using an automated high-content imaging system (Cellomics Array Scanner, VT1 HCS Reader, Thermo Fisher).

**Table 4.** Primary and secondary antibodies used for immunostaining.

| Antibody | Conc. | Host | Company | Catalogue# | RRID |
| --- | --- | --- | --- | --- | --- |
| Tau polyclonal Antibody | 1:500 | Rabbit | Santa Cruz Biotechnology | GG-GTX130462 | AB_2886280 |
| MAP2 polyclonal Antibody | 1:10,000 | Chicken | Abcam | ab5392 | AB_2138153 |
| NeuN | 1:500 | Rabbit | Merck | ABN78 | AB_10807945 |
| Tuj1 | 1:1,000 | Mouse | Promega Corporation | PRO-G7121 | AB_430874 |
| SSEA3 | 1:200 | Rabbit | DSHB | MC-631 IgM | AB_528476 |
| Alexa Fluor® 488 AffiniPure Donkey IgG | 1:200 | Rabbit | Jackson | 711-545-152 | AB_2313584 |
| Alexa Fluor® 488 AffiniPure Donkey IgG | 1:200 | Mouse | Jackson | 715-545-150 | AB_2340846 |
| Cy<sup>TM</sup>3 AffiniPure Donkey IgG | 1:200 | Chicken | Jackson | 703-165-155 | AB_2340363 |
| Alexa Fluor® 647 AffiniPure Donkey IgG | 1:200 | Rabbit | Jackson | 711-605-152 | AB_2492288 |

### High-content automated microscopy (HCA)

Automated microscopy of immunostained native or iNs was performed using a Cellomics Array Scanner (VT1 HCS Reader, Thermo Fisher) as previously described^33^. This platform enables rapid and unbiased image acquisition and analysis. Cell identification was carried out using the target activation (TA) protocol in CX5 HCS software. For each well of 24-well plates, 121 fields were captured with a 10× objective, and only wells containing at least 50 valid neurons were included in subsequent analyses.

The TA protocol defined DAPI^+^ cells based on fluorescence intensity, morphology, and area. Objects at image borders, aggregated nuclei, or unresolved clusters were excluded. Within the DAPI^+^ cells, TAU^+^, MAP2^+^, SSEA3^+^, NeuN^+^, and Tuj1^+^ cells were classified according to average cell body fluorescence intensity and area thresholds. iN culture purity was calculated as the proportion of marker-positive cells among total DAPI^+^ cells.

For neuronal morphology, each data point represents an individual donor cell line. Values reflect the mean of multiple single-cell measurements obtained from at least two independent wells (technical duplicates) per donor. Image acquisition and analysis of all wells were performed under identical HCA settings. Nuclei were considered valid based on the previously described criteria, with objects at the image borders excluded. Morphological analysis of TAU-stained cells - including cell body area, neurite number, neurite width, neurite length, neurite area, and branchpoint count - was quantified, and for each donor cell line, the reported values represent averages across all analyzed cells.

Thresholds for fluorescence intensity, area, and shape were established using randomly selected fields from both experimental groups. To ensure robustness, parameter settings were validated in a pre-analysis of 10 representative images per well and verified independently by a second researcher.

### Fluorescence-activated cell sorting (FACS)

To enrich for converted DPSC-derived neurons prior to RNA sequencing, cells were harvested and sorted by fluorescence-activated cell sorting (FACS) as previously described^34^. Cultures were dissociated with Accutase (Corning) at 37 °C for 10-15 min, washed in PBS, and centrifuged at 400 g for 5 min. After aspirating the supernatant, cells were resuspended in FACS buffer (HBSS supplemented with 1% bovine serum albumin (BSA) and 0.05% DNase I (Sigma)), centrifuged, and washed once more.

Cell suspensions were incubated with an allophycocyanin-conjugated antibody against human NCAM (1:10, PSA-NCAM-APC, Miltenyi Biotec, 130-120437, clone 2-2B) for 15 min on ice, protected from light. After antibody labeling, cells were washed twice in FACS buffer, centrifuged, and passed through a 70 µm cell strainer. Propidium iodide (1:1,000, Miltenyi Biotec, 130-093-233) was added immediately before sorting to exclude dead cells. Sorting was performed on a Beckman Coulter CytoFLEX SRT equipped with a 100 µm nozzle. Gates were set to collect NCAM⁺/PI⁻ live cells, and re-analysis confirmed sorting purity, with a cut-off of >80%.

Across experiments, the fraction of NCAM⁺ cells ranged from 11.6% to 30.6%, with post-sort purities of 82-97%. Between 9,700 and 26,600 NCAM⁺/PI⁻ single cells were obtained per sample. For downstream RNA-sequencing, sorted cells were collected at 10 °C directly into RLT buffer (Qiagen) containing 1% β-mercaptoethanol, snap-frozen on dry ice, and stored at −80°C.

### Quantitative real-time PCR (qRT-PCR)

Quantitative real-time PCR (qRT-PCR) was performed to measure RNA expression levels of neuronal, DPSC and pluripotency markers. Total RNA extraction was carried out using the RNeasy Mini or Micro Kit (Qiagen) according to the manufacturer’s instructions. cDNA synthesis was conducted using the Maxima First Strand cDNA Synthesis Kit. Primers (Table 5) were combined with LightCycler 480 SYBR Green I Master mix (Roche). The expression data were normalized using two reference genes - *ACTB* and *GAPDH*. Quantification was performed using the ΔΔCt metho

**Table 5.**
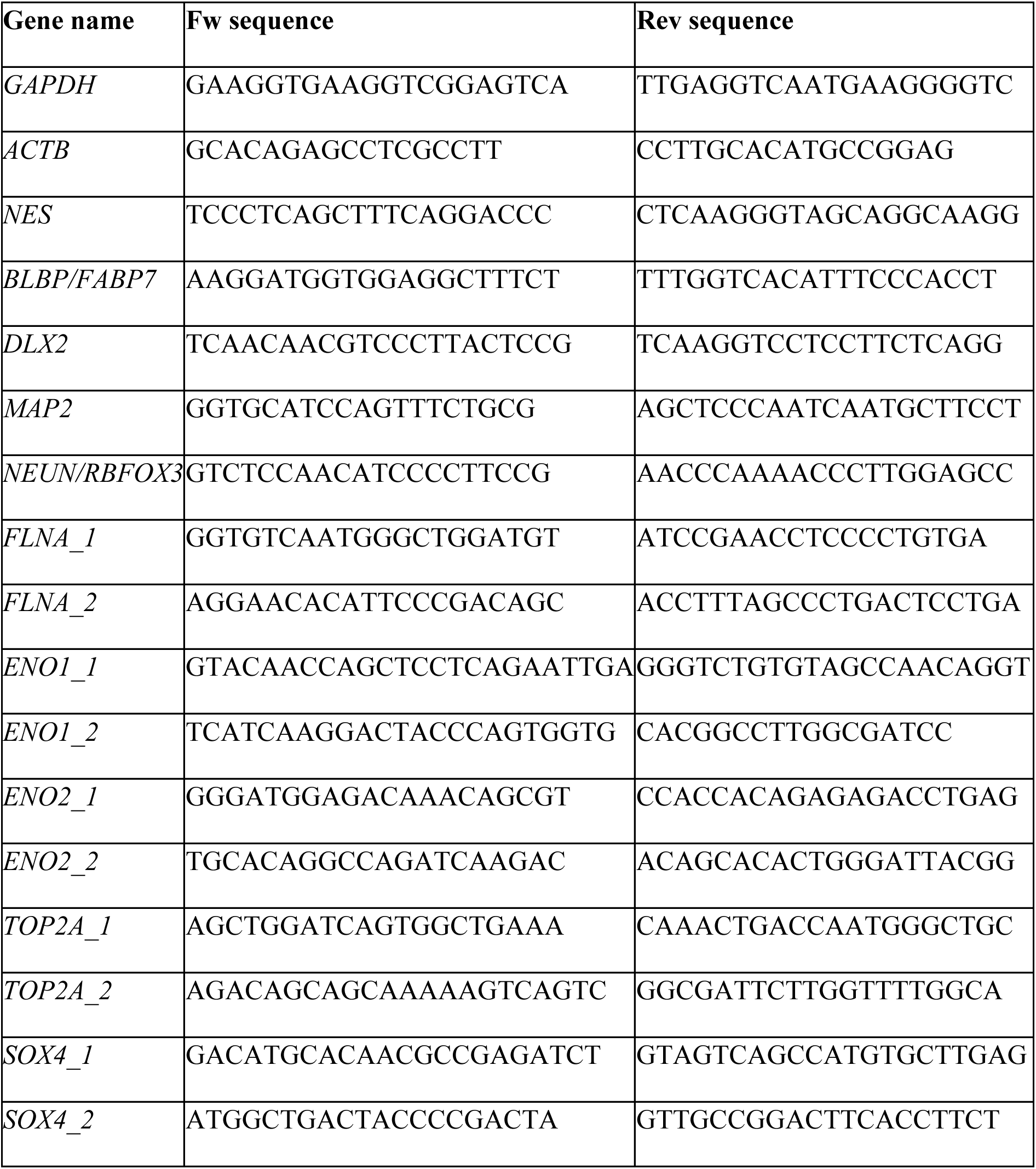
Primer sequences used for quantitative real-time PCR.

### Immunoblotting

For western blot (WB) experiments, 150,000 DPSCs were converted into iNs in a 0.1% gelatine-coated T25 flask from each sample. DPSC-iNs were harvested on day 17 by dissociating with Accutase (Corning) and collected with HBSS (Gibco) and spun at 400 g for 5 min. From native DPSCs 150,000 cells were pelleted and processed the same way. The cell pellets were lysed with RIPA buffer (Sigma) containing 4% cOmplete protease inhibitor cocktail (PIC) (Merck). Cell lysates were collected and incubated on ice for 30 min followed by centrifugation at 10,000 × g for 10 minutes at 4 °C to sediment cellular debris. The supernatant was collected and stored at −20 °C until further use. Gel electrophoresis and semi-dry blotting were performed as described previously^33^. Primary and secondary antibodies were diluted in 5% milk blocking solution (Table 6). For blot detection the Immobilon Western Chemiluminescent HRP Substrate (Millipore) was used, blots were incubated for 5 min at room temperature followed by immediate visualization by Imager2Imager CHEMI Premium Gel Documentation System (VWR).

**Table 6.** List of antibodies used for WB.

| Antibody | Conc. | Species | Company | Cat# | RRID |
| --- | --- | --- | --- | --- | --- |
| β-actin | 1:200,000 | Mouse | Sigma-Aldrich | A3854 | AB_262011 |
| Anti-βIII Tubulin/TUJ1 | 1:5,000 | Mouse | Promega | G7121 | AB_430874 |
| Hrp Conjugated antibody | 1:5,000 | Mouse | Cytiva | NA931 | AB_772210 |

### Whole-cell patch clamp electrophysiology

Whole-cell patch clamp recordings were performed on DPSC-iN cell cultures bathed in artificial cerebrospinal fluid (ACSF) consisting of: 140 mM NaCl, 5 mM KCl, 2 mM CaCl_2_, 1 mM MgCl_2_, 5 mM HEPES, and 10 mM D-glucose with pH = 7.5. Microelectrodes were pulled from 1B150F-4 borosilicate glass capillaries (World Precision Instruments) using a P-97 Flaming/Brown micropipette puller (Sutter Instruments) and then filled with intracellular solution consisting of 100 mM K-gluconate, 10 mM KCl, 20 mM KOH, 2 mM MgCl_2_, 2 mM NaCl, 10 mM HEPES, 0.2 mM EGTA, and 5 mM D-glucose. The electrode’s resistance was 4-6 MOhm, while the observed access resistance after membrane break-in was commonly less than 12 MOhm. During the experiments, cells were visualized using a Scientifica SliceScope microscope with an Olympus UMPLFLN 20X water immersion objective, and Qimaging digital camera with Qcapture software. Current clamp measurements were performed using a traditional current step protocol: 400 ms hyperpolarizing and depolarizing stepwise currents (from −30 pA to +50 pA with +2.5 pA increments) were injected to elicit voltage responses. When warranted by our observations on the current step responses (e.g. marked outward rectification or spikelets), voltage clamp recordings were performed to measure voltage-activated transmembrane currents. Here, cells were clamped at −60 mV holding potential and voltage steps were applied using 5 mV increments from −50 mV to +50 mV. All recordings were made using a MultiClamp 700B amplifier (Molecular Devices). To digitize the electrophysiological signals (20 kHz sampling frequency), we used a National Instruments PCIe-6220 data acquisition board with the DASYLab 2016 software in Windows 11. Recorded traces were analyzed using NeuroExpress (Szűcs, 2018; https://www.researchgate.net/publication/323547374_Analysis_of_miniature_excitatory_postsynaptic_currents_mini_analysis_in_NeuroExpress_Program_available_for_download).

### DNA methylation array

DNA methylation profiling was performed on both parental DPSCs and DPSC-derived iNs. Cells were cultured in T25 flasks, with 100,000 DPSCs seeded for differentiation, and genomic DNA was isolated using the DNeasy Blood and Tissue Kit (Qiagen) following the manufacturer’s instructions. Extracted DNA was subjected to bisulfite conversion with the EZ DNA Methylation Kit (Zymo Research). Genome-wide methylation status was then assessed using the Illumina Infinium MethylationEPIC v2.0 BeadChip (EPIC) array according to the supplier’s protocol.

### DNA methylation array data analysis

All data analysis was performed using R (v4.5.1)^35^. The IDAT files were imported, and beta values were computed using the openSesame pipeline from sesame package (v1.28.1)^36^. Several stages of filtering were performed to remove low quality probes. Firstly, probes with detection p-value > 0.01 and bead numbers < 3 in 10% of the samples were excluded. Further, non-CpG probes were removed, followed by SNP and cross-hybridising probes removal using rmSNPandCH function of the DMRcate package (v3.6.0)^37^.

M values of filtered probes were calculated using the BetaValueToMValue function of the sesame package, followed by annotation of the probe genomic coordinates using sesame’s EPICv2 hg38 Gencode v42 manifest file. Next, probes representing the same CpG site were aggregated by calculating the mean of the respective M values.

For promoter methylation analysis, promoters were defined as regions 1500 bp upstream and 500 bp downstream of Transcription Start Sites (TSS) of MANE select transcripts from MANE annotations (v1.4) of GRCh38 human transcriptome^38^. Filtered CpGs, with coordinates overlapping with that of the promoter regions were mapped to the respective genes. In cases of overlapping promoter regions, to avoid redundancy, CpGs within were mapped to genes with the closest TSS. Only promoter regions represented by at least 3 CpGs were further considered. M values for promoter regions were calculated by the average of assigned CpGs. Finally, differential methylation analysis of the promoter regions was performed using limma (v3.66.0)^39^ with a paired design for samples from the respective donors, followed by Benjamini-Hochberg correction for multiple testing. Significance was measured at adjusted p-value < 0.05.

### Epigenetic age prediction using DNA methylation clocks

Epigenetic age was predicted using the DNA Methylation Age Calculator of the Clock Foundation Team (https://dnamage.clockfoundation.org). We specifically tested the Horvath pan-tissue clock (*DNAmAge*)^40^, the skin and blood clock (*DNAmAgeSkinBloodClock*)^41^, and FitAge clock (*DNAmFitAge*)^42^. Additionally, we applied a methylation aging clock based on postmortem human cortical samples (*DNAmAge_cortical*)^43^.

### Bulk RNA sequencing

DPSCs (200,000 cells per sample) were expanded under standard culture conditions. For neuronal conversion, 500,000 cells were plated in T75 flasks and subjected to the established direct reprogramming protocol to generate induced neurons (iNs). Total RNA was extracted from 15 000 NCAM^+^/PI^-^ DPSC-iNs with the RNeasy micro kit (Qiagen) according to the manufacturer’s protocol. The sequencing libraries were synthesized with the SMART-Seq mRNA Kit (Takara/Clontech) and sequenced on a Novaseq X plus (paired end 2 × 150bp).

### Bulk RNA sequencing analysis

The raw sequencing data, available as .fastq files were mapped to the GRCh38.p14 human genome with Gencode hg38 annotations (v47) using the STAR aligner (v2.7.11b) tool with default settings^44^. Gene-level quantification was performed on the aligned and sorted bam files using the featureCounts function of the Subread (v2.0.8) tool with settings for paired-end reads and unstrandedness^45^.

All downstream analysis on the gene counts was performed using R (v4.5.1)^35^. The data were filtered to only include protein-coding genes with non-zero counts across all the samples. For principal component analysis, regularized log transformation was performed on the gene counts using rlog function from the DESeq2 package (v1.50.2)^46^, and the prcomp function from the stats package (v4.5.1) was applied on the top 500 most variable genes with “center” option set to TRUE. Differential expression analysis was performed using the DESeq2 package with a paired design for samples from the respective donors, followed by log-fold change shrinkage using the lfcshrink function with the ashr algorithm^47^. All genes with an adjusted p-value < 0.05 and absolute log2 fold-change > 0.5 were considered significantly differentially expressed. Functional enrichment analysis was performed using the clusterProfiler package (v4.18.3)^48^, separately for the significantly up- and downregulated genes. For over-representation test of gene ontology terms, the enrichGO function was used, while for that of reactome pathways, the enrichPathway function of the ReactomePA package (v1.54.0)^49^ was used. In both cases, significance was attributed at FDR < 0.05. Further, a semantic similarity-based clustering was performed on the significantly enriched gene ontology terms of up- and downregulated genes respectively, using the reduceSimMatrix function of the rrvgo package (v1.22.0), where similarity was computed based on the Relevance distance method and terms were reduced at a threshold set at 0.7^50^.

### Liquid Chromatography-Mass Spectrometry (LC-MS)

For LC-MS measurement, 300,000 DPSCs were seeded in T75 flasks for neuronal induction, and both the resulting DPSC-iNs and the corresponding native DPSC samples were collected and submitted for processing, with all pellets washed three times with PBS. Next, DPSC and DPSC-iN pellets were solubilized in 30 µl of 20% SDS then 6 µl of 1M TEAB (pH 8.5) was added. Cell lysis was performed by freeze-thaw sonication using a conventional ultrasonic bath. The protein concentration of the samples was determined by the BCA colorimetric assay, and sample aliquots representing 20 µg total protein were digested with trypsin using the suspension trapping workflow^51^. Briefly, protein disulfide bridges were reduced with TCEP (tris(2-carboxyethyl)phosphine), and the resulting free thiol groups blocked using MMTS (S-methyl methanethiosulfonate) and proteins were digested at 47 °C for 2h using 1 µg side-chain protected trypsin. The peptide samples were dried down and redissolved in 50 mM TEAB (pH 8.5). Sample aliquots representing 10 µg of original total protein were labeled with the TMTpro 16plex isobaric labeling tags according to the vendor’s protocol^52^. The labeled samples were mixed and dried down followed by high-pH reversed-phase fractionation using a C18 spin column according to the vendor’s protocol (Thermo Fisher Scientific, High-pH Reversed-Phase Peptide Fractionation Kit, User Guide MAN00084868). The collected fractions were dried down, redissolved in 0.1% formic acid (FA) and combined into four final fractions.

Fifty percent of the fractions were loaded onto disposable trap columns (Evotip) and analyzed with online LC-MS/MS using an Evosep One - FAIMS Pro - Orbitrap Fusion Lumos Tribrid mass spectrometer (Thermo Scientific) system. Peptides were separated using the „15 samples per day” built-in gradient (solvent A: 0.1% FA/water; solvent B: 0.1% FA/acetonitril; column: Evosep Endurance C18 Reprosil Pure, 0.15mm x 150mm ID x L, particle size: 1.9 µm, thermostated to 30 °C; flow rate: 220 nl/min). Mass spectrometric data acquisition was done in positive mode (emitter voltage: +2,300V) and in a data-dependent fashion. MS and MS/MS data were recorded at −50 and at −70 compensation voltages (CV) applying a 1.5 s cycle time for each CV. MS scans were recorded in the Orbitrap (resolution: 120,000, scan range: m/z 400 – 1,600, maximum ion inject time: 50ms, normalized AGC target: 100%). Multiply charged (z = 2-5) precursor ions were selected for MS/MS (intensity threshold: 50,000) with an isolation window of 0.7 Da. Dynamic exclusion time was 60s with 10ppm mass accuracy. MS/MS activation was HCD using 38% normalized collision energy. Fragment ions were detected in the Orbitrap (resolution:50,000, maximum ion inject time: 200ms, normalized AGC: 200%). Raw data were processed using Thermo Scientific Proteome Discoverer Software (v3.0, Thermo Fisher Scientific, San Jose, CA). Proteins were identified with the Sequest HT search engine (search parameters: database: SwissProt human sequences, downloaded 2024-10-02; enzyme: trypsin with a maximum of two missed cleavages; mass accuracy: 5 ppm for MS and 0.02 Da for MS/MS; modifications: fixed: methylthio on Cys, TMTpro on peptide N-termini and Lys; variable: Met oxidation, allowing up to two variable modifications per peptide). Peptide identifications with a minimum HT Xcorr value of 1 were retained, and protein identification confidence was set to “high”. Semiquantitative comparison was performed using the “unique + razor” peptide approach. Reporter-ion quantitation used signal-to-noise ratios (average S/N threshold: 10; precursor co-isolation threshold: 50%), and correction factors for TMTpro reagents were applied. Protein abundances were normalized to the total peptide amount.

All downstream data analysis was performed using R v4.5.1^35^. Proteins identified were first filtered to only include those identified with a confidence level marked as High. The normalized abundances of the filtered proteins were log2 transformed prior to further downstream steps. PCA was performed on the top 500 most variable proteins in the dataset using the prcomp function with “scale” and “center” options set to TRUE. Differential abundance analysis was performed using limma (v3.66.0)^39^ with a paired design for samples from the respective donors, followed by Benjamini-Hochberg correction for multiplicity. Proteins with an average log2 fold-change < −0.5 or > 0.5 and a corrected p-value < 0.05 were considered significant. An over-representation test of reactome pathways on the up- and downregulated proteins respectively, was performed using the enrichPathway function of the ReactomePA package (v1.54.0)^49^ with Benjamini-Hochberg correction and significance measured at FDR < 0.05.

### Nuclei extraction

For single nuclei isolation 300,000 cells were seeded into T75 flasks for neuronal conversion and pelleted and frozen at day 17. Cell suspensions were prepared from frozen cell pellets as previously described^53^. DPSC-iN samples were stored at −80 °C then thawed on ice until a homogeneous consistency was obtained. The cell pellets were transferred to a pre-chilled glass douncer in ice-cold lysis buffer (0.32 M sucrose, 5 mM CaCl₂, 3 mM MgAc, 0.1 mM Na₂EDTA, 10 mM Tris-HCl pH 8.0, freshly supplemented with 1 mM DTT and 0.1% Triton X-100; for RNA-seq experiments, 0.2 U/µL RNase inhibitor was added). Homogenization was performed with 10 strokes of the loose pestle followed by 10 strokes of the tight pestle.

The homogenates were transferred to 1.5 mL tubes and centrifuged at 500 × g for 5 minutes at 4 °C. The nuclear pellets were gently resuspended in ice-cold DPBS containing 2% BSA. Nuclei suspensions were passed through a 30 µm nylon cell strainer into precoated FACS collection tubes (PBS + 2% BSA). Before FACS sorting, DRAQ7 was added to stain nuclei, and single nuclei were sorted using a 100 µm nozzle on a BD FACS Aria cell sorter.

### Single-nuclei RNA sequencing

For transcriptomic profiling, single-nucleus suspensions were processed using the Chromium GEM-X Single Cell 3^’^ Library and Gel Bead Kit (10x Genomics, v3 chemistry) according to the manufacturer’s instructions. Nuclei were loaded onto the Chromium Single Cell GEM-X Chip together with the reagent master mix and partitioned into nanoliter-scale Gel Beads-in-Emulsion (GEMs) using the Chromium X Controller (10x Genomics). For donor DPSC2, 3 and 4, 20,000 nuclei from DPSC-iNs were loaded on the chip. Single-cell barcoding, reverse transcription, cDNA amplification, and library construction were performed according to the standard protocol provided by the manufacturer (10x Genomics; User Guide, Chromium GEM-X Single Cell 3′ Chip Kit Rev C).

### Single nuclei RNA sequencing analysis

Raw base calls were demultiplexed to obtain sample-specific fastq files, and reads were aligned to the GRCh38 genome assembly using the Cell Ranger pipeline (10x Genomics Cell Ranger 9.0.0) with default parameters and the include-introns option set to true ^54^.

The resulting matrix files were used for downstream analysis in R (v4.5.1)^35^ using the standard workflow of Seurat (v5.5.0)^55^. Nuclei with read counts between 3,000 and 60,000, gene counts between 1,500 and 10,000, and less than 5% mitochondrial transcripts were included for the analysis. Additionally, genes expressed in fewer than 10 cells were excluded from the dataset. After log-normalization and scaling, the data were integrated using the harmony package (v2.0.2)^56^, where individual samples were regressed out, and the first 20 principal components were chosen. Clusters resolved at a resolution of 0.6 were annotated based on known cell-type markers and those identified using the FindAllMarkers function of Seurat. Doublets and multiplets were predicted using scDblFinder (v1.24.10)^57^ and removed from further analysis. For cluster-specific marker identification, only protein-coding genes, excluding mitochondrial genes, were included. Marker genes for a given cluster were considered significant when Benjamini-Hochberg adjusted p-values were < 0.05, average log2 fold-change was > 0.5, the percentage of cells expressing the gene in the cluster (pct.1) was > 50%, and the percentage difference of cells expressing the gene between the cluster and the remaining cells (pct.1 - pct.2) was > 20%. Enrichment of biological processes corresponding to significant marker genes was tested with a gene over-representation test using the clusterProfiler package (v4.18.4)^48^, with Benjamini-Hochberg correction for multiplicity.

Lineage trajectories and pseudotime were computed using the slingshot package (v2.18.0)^58^, with “slingshot” as the distance method, the “Fibroblast-Like_1” cluster as the starting point, and the “GABAergic_Neurons” and “Unknown_Fate” clusters as the end points. These assignments were selected a priori for the computational analysis based on the gene expression profiles. Using the tradeSeq package (v1.24.0)^59^, a GAM model was fit on the computed pseudotimes and the top 2,000 highly variable genes previously identified in the Seurat workflow. The startVsEndTest and diffEndTest functions were then used to identify genes differentially expressed between the start and end points, or between the end points of the computed lineages. Genes were considered significant if Benjamini-Hochberg adjusted p-values were < 0.05, the Wald statistic was above the mean, and the log2 fold-change was > 0.5 or < −0.5. Finally, only protein-coding genes in the results were considered for further analysis. Enrichment of GO terms corresponding to significantly differentially expressed genes was tested with a gene over-representation test using the clusterProfiler package (v4.18.4)^48^, with Benjamini-Hochberg correction for multiplicity.

### Transcriptomic age prediction using single-cell ageing clocks

The single-cell inhibitory neuron-specific transcriptomic aging clock of Muralidharan et al.^60^ was applied to the single-nucleus RNA sequencing data. The processed gene expression data were used, and the gene names were mapped either directly to the clock features or through the GENCODE v32 release. Missing values were handled by average imputation, using the average expression of the missing genes in the clock’s training dataset^60^. The mean prediction of the five clock models resulting from cross-validation was calculated and used for evaluation.

Predicted age of cells in different clusters was compared by fitting a Mixed Linear Model (MLM) to the predicted age including age, sex and cluster as covariates and the donor ID as the grouping variable to account for having multiple samples from each donor. A cluster-cluster comparison was also performed by fitting an MLM to each pair of clusters. Mean predicted age of cells for each donor and cluster was calculated and compared as well by fitting an Ordinary Least Square (OLS) model to the predicted age including age, sex and cluster as covariates. Similarly as before, cluster-cluster comparison was also performed based on the mean predicted age by paired t-tests. In the MLM and OLS models, the p-value of the coefficient of the cluster variable determined the significance of the difference between the clusters.

The relation of pseudotime and predicted age in Lineage 1 and Lineage 2 was assessed by a MLM fitted to the predicted age using age, sex and pseudotime as independent variables. The significance of the relation was determined by the p-value of the coefficient of the pseudotime variable in this model.

In the statistical models, the clusters were coded as follows: 0: Fibroblast-Like_1, 1: Fibroblast-Like_2, 2: Fibroblast-Like_3, 3: Intermediary_1, 4: Intermediary_2, 5: Maturing_Neuron_1, 6: Maturing_Neuron_2, 7: GABAergic_Neurons, 8: Unknown_Fate.

GO enrichment analysis was performed with the positive and negative clock feature genes expressed in at least one cell in the iN data using the online GO Enrichment Analysis tool available at geneontology.org. Enrichment of GO biological process terms was tested with Fisher’s exact test with FDR correction for multiple testing.

### Statistical analysis

Statistical analyses were performed using GraphPad Prism. For comparisons between two groups, paired or unpaired *t*-tests were used as appropriate, depending on whether samples were matched by donor or derived from independent donor groups. Two-tailed tests were applied unless otherwise specified. When more than two groups were compared, one-way ANOVA followed by an appropriate multiple-comparisons test was used. Data normality was assessed using the Shapiro–Wilk test. Data are presented as individual donor values with mean ± SEM. Further statistical details are provided in the corresponding figure legends.

## RESULTS

### Efficient direct conversion of human dental pulp stem cells into induced neurons

We first established and optimized a direct conversion approach to generate iNs from DPSCs derived from surgically removed wisdom teeth (Fig. 1a, Table 1). Four independent donor-derived DPSC lines were transduced with our previously described all-in-one lentiviral vector - previously shown to induce direct neuronal conversion from fibroblasts within 28 days - and efficiently converted into neurons within 17 days^31,33,61–65^. Brightfield imaging revealed a progressive transition from fibroblast-like, spindle-shaped DPSCs at day 0 - 3 to neuronal-like morphologies by day 7 - 13, with extensive neurite outgrowth and complex arborization evident by day 17 (Fig. 1b). ICC-stained DPSC-iNs also confirmed that this neuronal morphology was reproducibly obtained from all four donor lines (Supplementary Fig. 1a).

To quantify changes in cellular identity during reprogramming, we performed HCA on unconverted parental DPSCs, and day-17 DPSC-iNs stained for neuronal and progenitor markers. Compared with native DPSCs, DPSC-iNs showed a significant reduction in the progenitor-associated marker SSEA3 and a marked increase in the fraction of DAPI⁺ cells expressing the neuronal markers NeuN, TUJ1, MAP2 and TAU (Fig. 1c, d). HCA also revealed a significantly higher number of DAPI⁺ nuclei in DPSC-iN cultures than in donor-matched parental DPSC cultures (Supplementary Fig. 1b). WB analysis further validated the neuronal conversion, showing significantly increased TUJ1 protein levels in DPSC-iNs relative to donor-matched DPSCs (Fig. 1e).

Quantitative morphological profiling using HCA-based profiling of TAU-positive cells further demonstrated that DPSC-iNs exhibited a consistent neuronal phenotype across donors, characterized by enlarged cell bodies, increased neurite number, longer neurite extensions, and higher branching complexity (Fig. 1f). HCA analysis showed that DPSCs gave rise to iNs with significantly higher purity compared with age- and sex-matched fibroblast-derived iNs (Supplementary Fig. 1c).

Finally, RT-qPCR analysis was consistent with the ICC results as DPSC-iNs showed increased expression of neurogenic and neuronal genes, including *NES*, *BLBP*/*FABP7*, *SOX4*, *DLX2*, *MAP2*, and *NEUN*/*RBFOX3*. In contrast, the mesenchymal/structural and metabolic markers *CD44*, *FLNA*, *ENO1*, and *ENO2* were reduced relative to parental DPSCs (Supplementary Fig. 1e). Together, these results demonstrate that human DPSCs can be rapidly and efficiently reprogrammed into iNs with high purity and neuronal morphology.

### Transcriptomic and proteomic profiling reveal neuronal identity in DPSC-iNs

To define the molecular changes associated with direct neuronal conversion, we performed bulk RNA sequencing and LC-MS/MS proteomics on parental DPSCs, and DPSC-iNs obtained from the four independent donors (Fig. 1a). Specifically, RNA sequencing was performed on NCAM1^+^-sorted DPSC-iNs. Upon performing principal component analysis (PCA) on the top 500 most variable protein-coding genes, we observed a strong distinction between DPSCs and DPSC-iNs at both the transcriptomic (PC1: 95% variance) and proteomic levels (PC1: 82% variance), indicating extensive molecular remodeling following conversion in multiple levels of gene expression (Fig. 2a). To identify the genes differentially expressed at the transcriptomic level, we performed a paired comparison on protein-coding gene expression across all the donors and identified 4,933 genes significantly upregulated while 3,766 genes downregulated in DPSC-iNs (NCAM1+) when compared to the respective parental DPSCs (Fig. 2b, Supplementary Data 1). Further, at the protein level, with a similar paired comparison, we found 1,223 proteins to be significantly upregulated in the DPSC-iNs, while 925 were downregulated (Fig. 2b and Supplementary Data 4).

**Figure 2:**
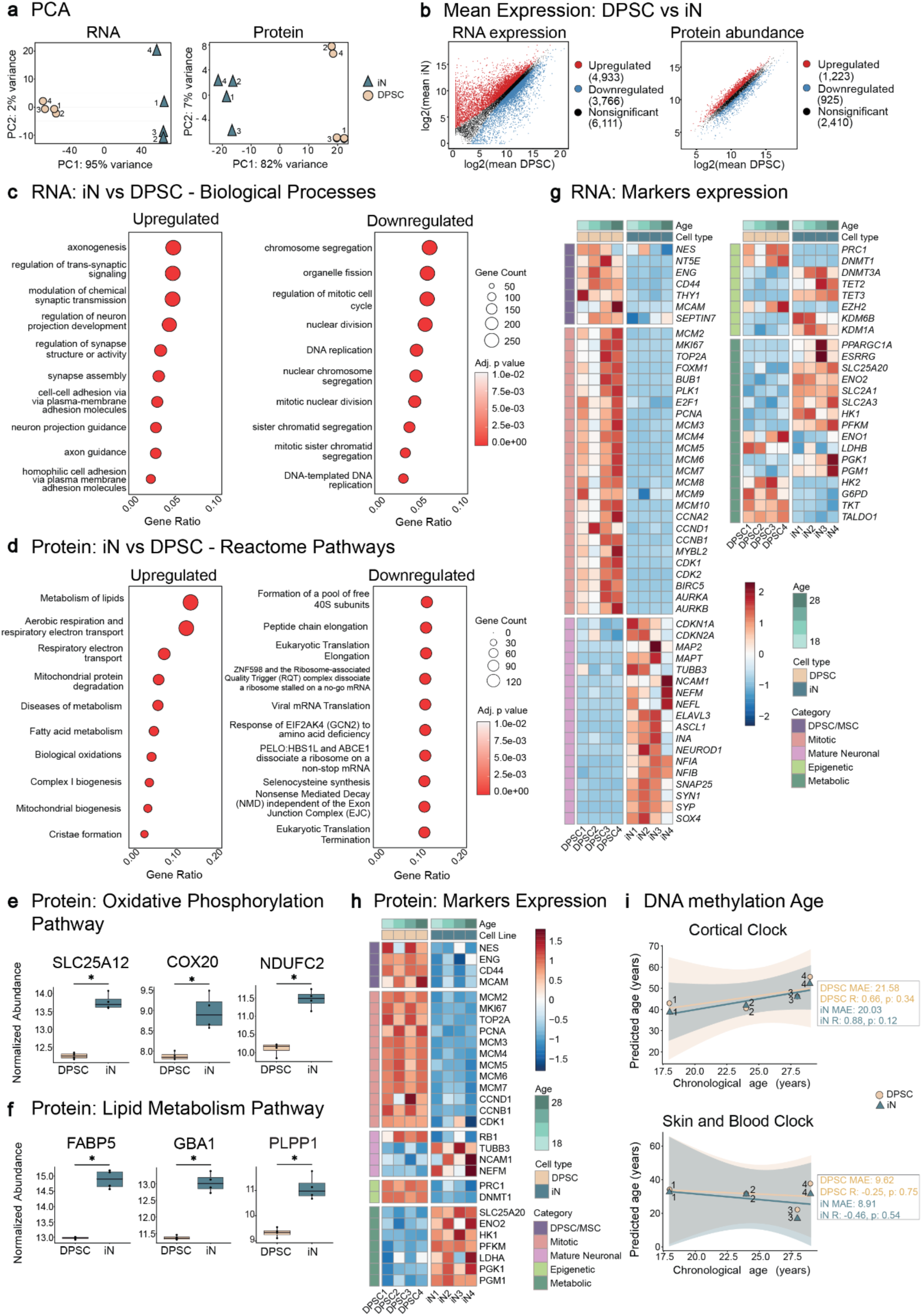
Multi-omic profiling reveals neuronal identity of DPSC-iNs. **a:** Plots of principal components from PCA of top 500 (left) protein-coding genes from bulk RNA-seq and (right) proteins from MS-based proteomics datasets showing clear separation between DPSCs and DPSC-iNs^#^ across four donors (n = 4 donors). **b:** Mean expression levels of (left) protein-coding genes derived from bulk RNA-seq and (right) proteins derived from MS-based proteomics data compared between DPSCs and DPSC-iNs^#^ (n = 4 donors). **c:** Biological processes enriched for genes (left) upregulated and (right) downregulated in DPSC-iNs^#^ when compared to DPSCs in the bulk RNA-seq data. **d:** Reactome pathways enriched for proteins (left) upregulated and (right) downregulated in DPSC-iNs when compared to DPSCs in the MS-based proteomics data. **e:** Normalized abundance of example oxidative phosphorylation proteins derived from MS-based proteomics data and significantly increased in DPSC-iNs (n = 4 donors). **f:** Normalized abundance of example lipid-metabolism proteins derived from MS-based proteomics data and significantly increased in DPSC-iNs (n = 4 donors). **g, h:** Heatmaps showing (g) normalized transcriptomic expression of selected differentially expressed genes and (h) normalized abundance of selected differentially abundant proteins characterizing the identity of the DPSC and DPSC-iN^#^ cell lines. Statistical significance was determined using ClusterProfiler in c and d, DESeq2 in b (left) and g, and limma in b (right), e, f and h, coupled with Benjamini-Hochberg correction. **i:** Plot showing correlation of chronological age with predicted DNA methylation age based on (top) Cortical clock and (bottom) Skin and Blood clock in the DNA methylation array data of DPSCs and DPSC-iNs (n = 4 donors). Using Pearson’s correlation test, the correlation coefficient R, and p-value were determined separately for the DPSCs and DPSC-iNs. ^#^ For bulk RNA-sequencing, DPSC-iNs were FACS sorted to only include NCAM1+ cells (Refer Methods). * Significance at adjusted p-value < 0.05.

When we performed a functional enrichment analysis on the genes or proteins differentially expressed, we found further evidence highlighting the shift in molecular profile between DPSCs and DPSC-iNs. Using a gene ontology analysis on the genes upregulated in DPSC-iNs (NCAM1+) at the RNA level, we found an enrichment of biological process related to axonogenesis and synaptic signaling (Fig. 2c, Supplementary Fig 2a and Supplementary Data 2). Conversely, the downregulated genes were associated with processes related to DNA replication and mitosis (Fig. 2c, Supplementary Fig 2a and Supplementary Data 2). Overall, this suggested the enrichment of neuronal processes and the reduction of mitotic and mesenchymal programs at the transcriptomic level. We observed similar trends at the proteomic level where pathway enrichment analysis, with the Reactome database, associated upregulated proteins with neuronal metabolic pathways, such as aerobic respiration, oxidative phosphorylation and lipid metabolism (Fig. 2d and Supplementary Data 5). Likewise, the downregulated proteins were associated with RNA processing and mitotic cell-cycle pathways (Fig. 2d and Supplementary Data 5). Furthermore, important proteins belonging to the oxidative phosphorylation pathway, such as SLC25A12, COX20, NDUFC2 and those belonging to the lipid metabolism pathway, such as FABP5, GBA1 and PLPP1, known to be associated with neuronal maturation program, were significantly upregulated in DPSC-iNs (Fig. 2e, f, Supplementary Fig. 2b, 2c and Supplementary Data 4).

Upon checking the expression of markers to validate cellular identity, we found several to be significantly differentially expressed, confirming the global transcriptomic and proteomic shifts from a mesenchymal stem cell to a neuronal identity. At the RNA level, DPSCs showed higher expression of mesenchymal stem cell markers (*ENG*, *MCAM, THY1*, *CD44*), proliferative markers (*MKI67*, *TOP2A*, *PCNA*, multiple MCM family members), and epigenetic regulators characteristic of stem-like states (Fig. 2g). These markers were significantly downregulated in DPSC-iNs (NCAM1+) (Supplementary Data 1). Conversely, mature neuronal genes - including *MAP2*, *MAPT*, *TUBB3*, *NEFM*, *SNAP25* and *SYN1* - were upregulated in the iNs (Fig. 2g). We observed the proteomic data to mirror these results, where there was an increased abundance of neuronal proteins (NCAM1, NEFM, TUBB3, NEFL) and reduced abundance of proliferative markers (MCM2-7, PCNA, CCNB1, CDK1) in the iNs (Fig. 2h and Supplementary Data 4).

Overall, we identified substantial concordance in the changes across the transcriptomic and proteomic levels, largely in the same direction, with 4,326 genes uniquely upregulated and 3,501 uniquely downregulated at the transcriptomic level (Supplementary Fig. 2d, Supplementary Data 1 and 4). The concordant overlap indicated that many of the major transcriptional changes during reprogramming were also reflected at the proteomic level.

Together, our results show that DPSC-iNs undergo a coordinated transcriptional and proteomic transition characterized by the suppression of mesenchymal-like stem cell and mitotic states, and the activation of neuronal pathways and post-mitotic state, consistent with a neuronal identity, confirming the direct neuronal reprogramming of dental pulp stem cells.

### DNA methylation profiling reveals promoter-level remodeling during direct neuronal conversion

To assess whether direct neuronal conversion alters donor-associated epigenetic age in DPSC-iNs, we profiled parental DPSCs and matched day-17 DPSC-iN cultures using the EPICv2 DNA methylation array. We applied several DNA methylation aging clocks trained on cortical, skin and blood, or multiple tissue types (Fig. 2i, Supplementary Fig. 3a, and Supplementary Data 7). Although predicted methylation ages did not correlate significantly with chronological age in this small cohort, no significant differences were detected between matched DPSCs and DPSC-iNs (Supplementary Data 7). This pattern is consistent with relative stability of methylation-age estimates during conversion, although it does not by itself establish preservation of donor-specific epigenetic aging signatures.

We next examined conversion-associated methylation changes at protein-coding gene promoters. A principal component analysis on the top 500 promoters revealed strong donor-donor differences (Supplementary Fig. 3b). By performing a differential methylation analysis on the protein-coding gene promoters, we identified promoters of 36 genes to be significantly hypermethylated in DPSC-iNs, while that of 1 gene to be hypomethylated (Supplementary Fig. 3c, d and Supplementary Table 6). Given the established role of promoter DNA methylation in regulating gene transcription, we checked whether the genes associated with differentially methylated promoters also exhibited altered transcriptomic expression (Supplementary Fig. 3d, Supplementary Data 1 and 6). The only identified hypomethylated gene in the DPSC-iNs, namely, phosphoglycerate kinase 1 (*PGK1*) was significantly upregulated (Supplementary Fig. 3d, Supplementary Data 1 and 6). *PGK1* encodes an important glycolytic enzyme implicated in axonal ATP production in neurons^66^. Additionally, only a few of the hypermethylated genes were found to be downregulated in DPSC-iNs, and these were associated with non-neuronal - epithelial, chondrogenic, or osteogenic lineages (Supplementary Fig. 3d, Supplementary Data 1 and 6).

Together, these findings suggest that direct conversion does not produce a detectable shift in methylation-age estimates between matched DPSCs and DPSC-iNs, while selectively remodeling promoter methylation at loci associated with metabolic and non-neuronal lineage programs. Although consistent with retention of donor-associated methylation features, studies involving larger cohorts and a broader donor-age range will be required to determine whether donor-specific epigenetic aging signatures are preserved during DPSC-to-iN conversion.

### DPSC-iNs exhibit developing neuronal electrophysiological properties

To assess functional maturation of DPSC-iNs, we performed whole-cell patch-clamp recordings in current-clamp configurations between day 14 - 17 of neuronal conversion (Fig. 3a). Whole-cell recordings were obtained from 23 DPSC-iN cultures across four donors (Fig. 3a-f; n = 98 cells in total). The recorded “differentiated” DPSC-iNs (n = 56) exhibited substantial heterogeneity in resting membrane potential and input resistance. However, the experiments also demonstrated that many of these cells exhibited features in their membrane potential that was consistent with the action of voltage-gated ion channels, a hallmark of neurons. These cells were classified as “responsive cells” (26 of 56) and comprised 46% of recorded differentiated DPSC-iN cells (Fig. 3c-f). This group exhibited signatures of voltage-sensitive ion channel activity within the physiological membrane potential range (from −95 mV to +20 mV). We observed a robust decrease in cell input resistance during depolarizing steps and action potential spikelets with an amplitude of ≤ 20 mV and a slow time course. Such high-resistance cells and slowly developing active responses are consistent with an immature neuronal electrophysiological state^67^.

**Figure 3.**
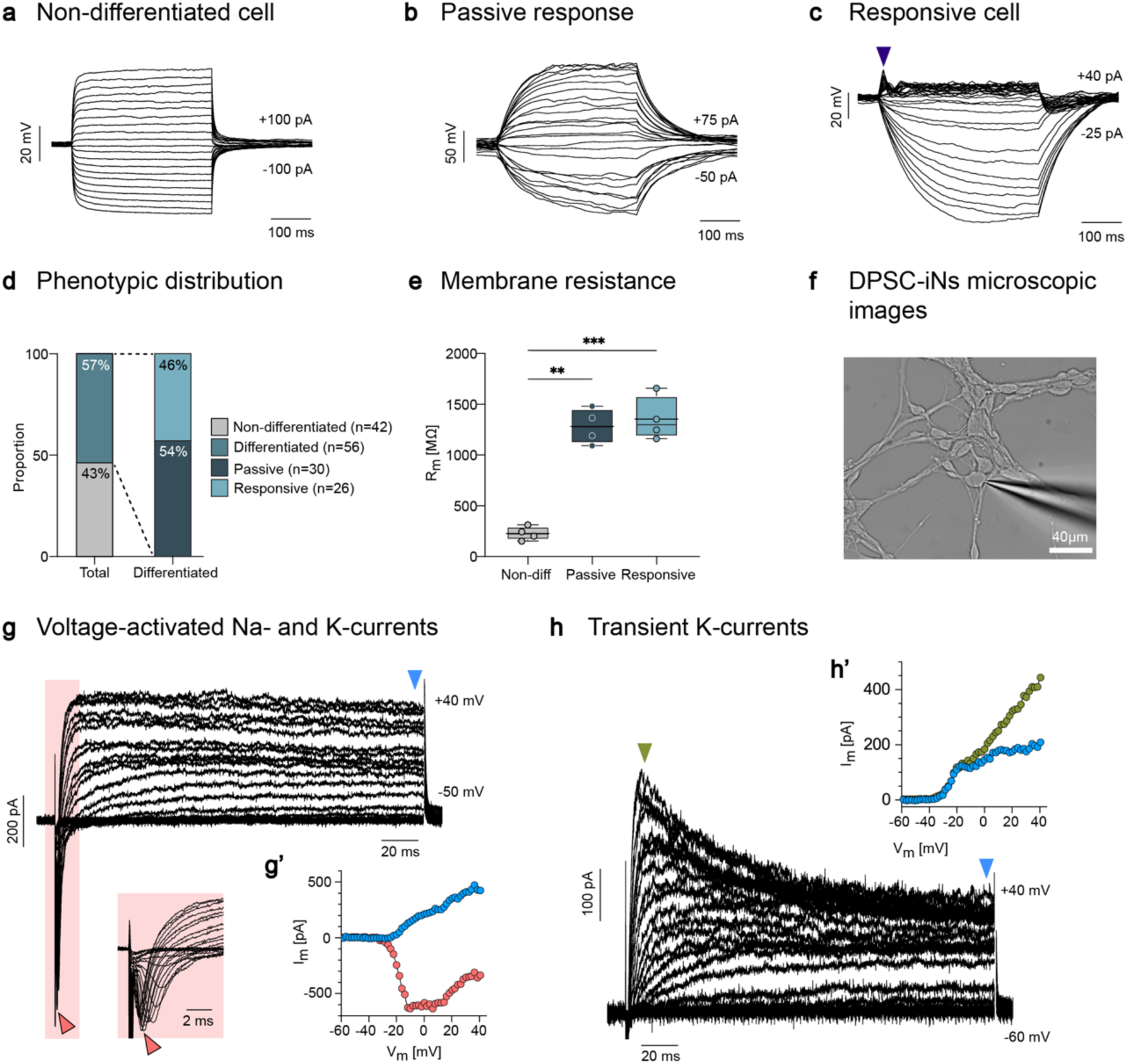
DPSC-iNs develop early neuronal electrophysiological properties. **a-c:** Representative whole-cell current-clamp recordings. **a:** Non-differentiated DPSCs exhibit low membrane resistance and membrane time constants, with no evidence of voltage-sensitive ion channel activity. **b:** A subset of the DPSC-iNs show passive properties, with large Vm changes indicating a low density of voltage-dependent ion channels in the cell membrane. **c:** A large percentage of DPSC-iNs were responsive cells that generated action potential spikelets in response to depolarizing steps, indicating the presence of active Na⁺ currents (arrow). Additionally, the cells exhibited outward rectification under positive input currents, as indicated by sublinear Vm development with incremental current strength. **d:** Proportion of all recorded cells across four donors (n = 98: 42 non-differentiated DPSCs, 30 passive DPSC-iNs, and 26 responsive DPSC-iNs). **e:** Membrane resistance values for each group. Boxes indicate the median and interquartile range; whiskers indicate the full range; individual points represent average membrane resistance values within each donors (n = 4 per group). **f:** Representative image of a recorded DPSC-iN.(scale bar: 40 µm) **g:** Voltage clamp recording of a responsive DPSC-iN reveals fast inward currents followed by sustained outward currents. **g’:** Current-voltage relationships for inward (red) and sustained outward (blue) components. (see corresponding arrows). **h:** Additional responsive cell showing prominent transient outward currents consistent with A-type K⁺ channels. **h’**: Corresponding current–voltage relationships. Statistical differences in were assessed by using one-way ANOVA followed by Tukey’s multiple-comparisons test. Significance: *p < 0.05, **p < 0.01, ***p < 0.001.

Meanwhile, 54% of recorded DPSC-iNs (30 of 56) were classified as “passive” DPSC-iNs with a high input resistance (>1000 MΩ), slow apparent membrane time constants (>20 ms), and a linear relationship of membrane potential amplitude to incremental strength input current steps (Fig. 3b, d and e). These characteristics resemble those of immature neuronal precursors typically observed in iPSC-derived neurons^68^, or fibroblast-iNs at early maturation stages^69^. By contrast, “non-differentiated” DPSCs (n = 42, 43% of the total) exhibited no neuronal electrical features and displayed low membrane resistance (<350 MΩ), rapid membrane time constants (<10 ms), and linear current-voltage relationships (Fig. 3a, d and e). Fig. 3a-c shows whole-cell recordings in undifferentiated cells and DPSC-iNs associated with the passive or responsive electrical phenotypes.

Follow-up voltage-clamp recordings in “responsive” DPSC-iNs confirmed the presence of inward and outward voltage-gated currents characteristic of developing neurons (Fig. 3g’). Transient inward currents, consistent with Na⁺ channel activity, are the drivers of spikelets seen in current clamp (Fig. 3c). Outward rectification was mediated by various types of K⁺ currents: slowly activating currents consistent with M-type channels (Kv7), and fast-activating and slowly inactivating A- or D-type K⁺ currents (Fig. 3h’).

Collectively, these results indicate that a subset of DPSC-iNs develops active electrophysiological properties within 17 days of conversion, displaying early-stage excitability and a repertoire of voltage-gated currents characteristic of immature but functionally developing neurons.

### Single-nucleus transcriptomics reveals GABAergic-like transcriptional fate in DPSC-iN cultures

Given that DPSCs are primary cells with an intrinsically heterogeneous cellular composition, and the observed heterogeneity in electrophysiological characterization of DPSC-iNs, we performed single-nucleus RNA sequencing to resolve the cellular diversity of DPSC-derived iN cultures after 17 days of direct neuronal reprogramming. This approach allowed us to assess whether distinct neuronal maturation states or non-neuronal fates could be detected within the differentiated cultures.

Single-nucleus RNA sequencing was performed across three DPSC-iN samples. After sequencing and quality-control (Supplementary Fig. 4a-d), we obtained 19,418 high-quality nuclei distributed across 9 clusters (Fig. 4a). Based on the top genes expressed in each cluster (Supplementary Data 8), and established cell type markers (Fig. 4b), we identified fibroblast-like cells, neuronal clusters, and several transitional cell states that co-expressed neuronal and mesenchymal lineage genes (Fig. 4b-e and Supplementary Fig. 4e). Among the neuronal clusters, the Maturing Neurons 1 and 2 clusters showed high expression of immature neuronal markers, including *STMN3* and *TUBB3*, together with the Maturing Neurons 2 cluster showing lower expression of mature neuronal markers such as *MAP2* and *MAPT*, consistent with intermediate stages of neuronal maturation (Fig. 4b and Supplementary Data 8). In contrast, the GABAergic Neurons cluster showed higher expression of mature neuronal markers together with GABAergic or interneuron-associated genes, including *DLX5* and *LHX8* (Fig. 4b and Supplementary Data 8). Interestingly, the GABAergic neuron cluster also showed expression of immature neuronal genes, suggesting that these cells might still be in the process of maturation at day 17 of conversion. (Fig. 4b and Supplementary Data 8). We annotated an additional cluster as Unknown Fate as it lacked a clear neuronal gene expression profile while retaining expression of some fibroblast associated markers (Fig. 4b and Supplementary Data 8). None of the identified clusters showed evidence of active proliferation, as indicated by a lack of *TOP2A* and *MKI67* expression (Fig. 4b and Supplementary Data 8), further supporting the post-mitotic status observed in the bulk RNAseq and proteomic data of DPSC-iNs.

**Figure 4.**
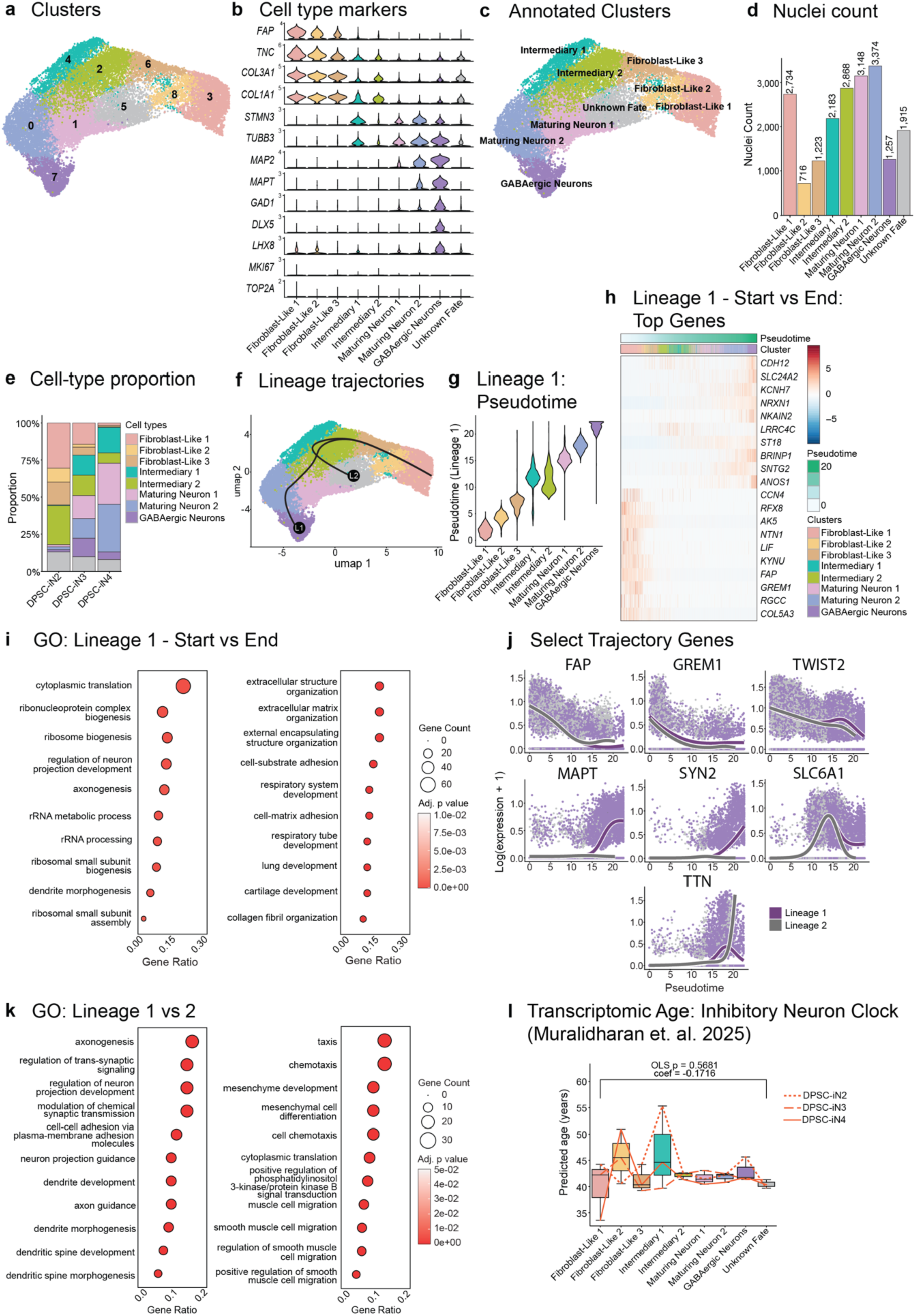
Single-nucleus transcriptomics identifies heterogeneous states in DPSC-derived iN cultures. **a:** UMAP plot of clusters identified in the day 17 DPSC-iN snRNA-seq data (n = 3). b: Violin plots showing the expression levels of selected genes characterizing the identity of the clusters in the day 17 DPSC-iN snRNA-seq data (n = 3). c: UMAP plot of clusters annotated by cell type or fate in the day 17 DPSC-iN snRNA-seq data (n = 3). d: Number of high-quality cells identified in each cluster in the day 17 DPSC-iN snRNA-seq data (n = 3). e: Proportion of cells from each cluster in every donor in the day 17 DPSC-iN snRNA-seq data (n = 3). f: UMAP plot showing the predicted pseudotime-based lineage trajectories, lineage 1 (L1) and lineage 2 (L2) in the day 17 DPSC-iN snRNA-seq data (n = 3). g: Violin plots showing computed pseudotime values per cluster in lineage 1 of the day 17 DPSC-iN snRNA-seq data (n = 3). h: Heatmap showing the normalized and scaled expression of top 10 genes differentially expressed in the start or end points of the lineage 1 trajectory in the day 17 DPSC-iN snRNA-seq data (n = 3). i: Top biological processes enriched for genes differentially expressed between the start and end points of lineage 1 trajectory in the day 17 DPSC-iN snRNA-seq data (n = 3). j: Plots showing trend of log2 normalized expression of selected genes against predicted pseudotime each in lineage 1 and lineage 2 trajectories in the day 17 DPSC-iN snRNA-seq data (n = 3). k: Top biological processes enriched for genes differentially expressed between the end points of lineage 1 and lineage 2 trajectories in the day 17 DPSC-iN snRNA-seq data (n = 3). l: Donor-level cluster-wise average of transcriptomic ages predicted by the Single-cell Inhibitory Neuron clock from Muralidharan et al. 2025, in the day 17 DPSC-iN snRNA-seq data (n = 3). An ordinary least squares model was fit on the predicted age and chronological age, with cluster and sex as additional variables, and significance was measured at p-value < 0.05. Statistical significance was determined in h and j, using tradeseq’s Wald test, and in i and k using ClusterProfiler, both coupled with Benjamini-Hochberg correction. False-discovery was controlled at corrected p-value < 0.05.

Given the presence of various transitional and putative terminal cell states, we next computationally inferred lineage trajectories using Slingshot, with the Fibroblast-Like 1 cluster specified as the starting state and the GABAergic Neurons and Unknown Fate clusters specified as endpoints (Fig. 4f). The analysis identified two major lineage trajectories (Fig. 4f). Lineage 1 connected fibroblast-like and intermediary states with the maturing neuronal clusters and the GABAergic Neurons cluster (Fig. 4f-h). Lineage 2 extended from the Fibroblast-Like 1 cluster through Intermediary 1 toward the Unknown Fate cluster (Supplementary Fig. 4f). Along Lineage 1, neuronal genes such as *BRINP1* and *NRXN1* showed increased expression with pseudotime, while mesenchymal genes such as *FAP*, *COL5A3* and *GREM1*, were decreased (Fig. 4j and Supplementary Data 9). Consistent with these expression patterns, genes increasing along Lineage 1 were enriched for processes related to axonogenesis and synapse formation, whereas genes decreasing along the trajectory were enriched for extracellular matrix organization (Fig. 4i and Supplementary Data 10). Along Lineage 2, which shared the prespecified starting cluster Fibroblast-Like 1, genes increasing in expression with pseudotime were enriched for processes related to smooth-muscle cell migration, suggesting an alternative non-neuronal transcriptional state (Supplementary Fig. 4f-h and Supplementary Data 9). Comparison of the two lineage trajectories showed a shared decrease in mesenchymal-associated genes expression, including *FAP*, *GREM1*, and *TWIST2*, with increasing pseudotime (Fig. 4j, k, Supplementary Fig. 4i, and Supplementary Data 9). By contrast, the putative neuronal lineage showed a distinct increase in the expression of neuronal genes, including *MAPT*, *SYN2*, and *SLC6A1* (Fig. 4j, k, Supplementary Fig. 4i and Supplementary Data 9). Genes preferentially associated with the alternative endpoint were enriched for smooth-muscle-related biological processes (Fig. 4k and Supplementary Data 10). Among the genes enriched at the Unknown Fate endpoint were developmental and Wnt/BMP-associated genes, such as *LEF1*, *NKD1* and *BMP7*, extracellular-matrix-associated genes, including *POSTN*, *LOXL2*, *TGFBI* and *HAS2*, and mechanosensory-associated genes, including *PIEZO2* and *TRPM3*. This expression profile is compatible with a dental mesenchymal or odontoblast-lineage precursor-like state. However, several neuronal-development-associated genes also increased along Lineage 2, and the cluster did not exhibit a definitive mature odontoblast or sensory-neuron signature. We therefore interpret the Unknown Fate cluster as a putative transitional neural crest-derived state rather than a fully committed lineage.

To determine whether neuronal maturation was accompanied by measurable differences in transcriptomic age, we applied our previously developed single-nuclei transcriptomic aging clocks to the DPSC-iN clusters^60^. Particularly, given that the data showed a commitment to an inhibitory neuronal fate, we applied the single-cell inhibitory neuron ageing clock. This enabled cell-level estimation of transcriptomic age across mature and less mature neuronal populations, as well as across transitional and non-neuronal cell states. To test the applicability of the clock in the dataset, we first checked the number of clock features expressed in the dataset and identified 62.5% of the features to be expressed in at least one cell (Supplementary Fig. 5a and b). Further, among the overlapping features, the genes with a negative coefficient were associated with biological processes related to neurodevelopment, while those with a positive coefficient were associated with metabolic processes (Supplementary Fig. 5c and Supplementary Data 11), further highlighting their relevance. Overall, we observed no significant differences in the average transcriptomic age between the clusters regardless of the maturation status or the cell fate (Fig. 4l, Supplementary Fig. 5d and Supplementary Data 12). Further, each cluster had 82-85% of the cells with predicted age above the chronological age, while 8-10% of the cells had a negative predicted age (Supplementary Fig. 5e and Supplementary Data 12). Additionally, we detected no significant association between predicted transcriptomic age and pseudotime along either of the inferred branches (Supplementary Fig. 5f and g).

Together, these results indicate that day-17 DPSC-iN cultures contain heterogeneous conversion states, including maturing neuronal, GABAergic, fibroblast-like, intermediary, and alternative non-neuronal populations. The computationally inferred lineages were indicative of two alternate fates, neuronal and non-neuronal endpoints. However, given that the analysis was performed at a single time point of day 17, the inferred lineages require further experimental validation to establish their relationships and directionality. Finally, the exploratory application of the clock enabled us to speculate that the transcriptomic age remains unchanged during potential cell-state transitions.

## DISCUSSION

In this study, we provide an integrated characterization of rapid, transcription factor-mediated direct neuronal conversion of human DPSCs. By combining transcriptomic, proteomic, DNA methylation, single-nucleus RNA-sequencing, and electrophysiological analyses, we show that DPSCs can be converted within 17 days into iNs that acquire neuronal molecular features and early electrophysiological properties. Compared with fibroblast-derived iNs generated using the same reprogramming construct under the respective established protocols, DPSC-iNs showed accelerated conversion kinetics (17 versus 28 days) and higher neuronal purity, suggesting the developmental advantages of neural crest-derived donor cells for neuronal reprogramming.

The multi-omic investigation revealed extensive transcriptomic and proteomic shifts during conversion, including the downregulation of genes associated with mesenchymal stem cell identity and the mitotic cell cycle, together with the activation of neuronal programs. In particular, DPSC-iNs showed expression of neuronal markers and genes critical for synaptic signaling, oxidative phosphorylation, and lipid metabolism, consistent with the metabolic and functional shifts associated with a post-mitotic neuronal state^70^.

DNA methylation-clock estimates did not differ significantly between matched DPSCs and DPSC-iNs, suggesting that the donor age-related methylation patterns might be preserved. However, since the predicted ages did not significantly correlate with the chronological age, possibly limited by the small cohort and a narrow donor-age range, this needs to be further explored in a larger cohort. Determining whether DPSC-iNs retain donor-associated molecular age remains important for their potential application in modeling late-onset neurodegenerative diseases, particularly because reprogramming through pluripotency can reset age-associated features^6,7^.

Electrophysiological profiling provided evidence of early neuronal functional development. Within 17 days, a subset of DPSC-iNs developed active membrane properties, including spikelet generation and the expression of voltage-gated Na⁺ and K⁺ currents. Although these properties reflect an immature stage, they support the acquisition of early neuronal electrophysiological features and align with early stages observed in iPSC- and fibroblast-derived iNs^68,69^. Importantly, functional maturation may be enhanced by culturing DPSC-iNs on an astrocyte feeder layer, an established strategy to promote synaptogenesis and excitability in developing neuronal cultures^71,72^. Thus, while early intrinsic activity was observed in our system, the inclusion of astrocytic support may facilitate more advanced functional outcomes in future work.

A key obstacle in translating DPSCs into reproducible cell platforms lies in their heterogeneity and limited proliferative capacity as primary cells. To address this, we performed single-nucleus RNA sequencing of day-17 DPSC-iN cultures, enabling us to resolve the cellular composition generated during direct reprogramming. We found that DPSC-to-iN conversion at day 17 does not include a uniform neuronal population, but contains multiple maturation states, including maturing neuronal clusters and a GABAergic-like neuronal cluster, alongside residual non-neuronal or alternative-fate populations such as the “Unknown Fate” cluster. Lineage trajectory inference further predicted a putative neuronal maturation path toward a GABAergic neuronal fate and a separate path toward the Unknown Fate cluster. The mixed dental mesenchymal/odontoblast-lineage precursor-like and neuronal-development-associated expression features of this cluster are compatible with a transitional neural crest-derived state rather than a fully committed lineage.

Additionally, upon application of the single-cell inhibitory-neuron transcriptomic aging clock described by Muralidharan et al. (2025)^60^, we observed no significant differences in predicted transcriptomic age across cell populations or along either inferred lineage, despite differences in maturation state and the expression of immature neuronal markers among several clusters. Notably, a substantial proportion of the clock features was represented in the DPSC-iN dataset, including genes associated with neuronal and age-related biological processes, supporting the relevance of these cells to inhibitory-neuron aging programs. However, given that the clock was developed based on adult postmortem brain samples and our donors represented a young and relatively narrow age range, the resulting predictions should not be interpreted as precise estimates of biological age. Together, these analyses indicate the potential conversion paths and the cell fates acquired during direct neuronal conversion of DPSCs. Future lineage-tracing studies will be required to validate the inferred trajectories and determine whether specific starting DPSC subpopulations preferentially contribute to neuronal or GABAergic-like states. Furthermore, enrichment or selection of such favorable DPSC subpopulations may improve conversion purity and extend the usable passage window for direct reprogramming applications.

Taken together, our findings support DPSCs as a developmentally relevant and readily accessible donor cell source for direct neuronal conversion. By integrating multi-omics, electrophysiological profiling, single-nucleus transcriptomics, trajectory inference, and transcriptomic age prediction, we provide the first framework for understanding the molecular, cellular, and functional basis of DPSC-to-iN direct conversion. Beyond advancing basic knowledge, this work highlights the potential utility of DPSCs for patient-specific disease modeling, particularly for studies that benefit from rapid conversion and the direct use of patient-derived somatic cells. Because this study included only four donors within a narrow age range, larger cohorts spanning a broader range of ages will be required to determine whether DPSC-to-iN conversion preserves donor-associated aging signatures. Importantly, the availability of DPSCs not only from surgically removed wisdom teeth but also from naturally exfoliated deciduous “milk” teeth opens the possibility of generating patient-specific neuronal models of pediatric disorders. Future efforts incorporating astrocytic co-culture and promoting long-term maturation will further enhance the utility of DPSC-iNs for neurobiology and disease modeling.

## DATA AVAILABILITY

The bulk RNA-seq, single-nucleus RNA-seq, and DNA methylation datasets generated in this study have been deposited in the Gene Expression Omnibus and will be made publicly available upon peer-reviewed publication. The mass spectrometry dataset has been deposited in PRIDE and will be made publicly available upon peer-reviewed publication.

## CODE AVAILABILITY

All custom code and software pipelines developed for data preprocessing, formal analysis, and visualization are publicly available on GitHub, at https://github.com/chanmur/DPSC_iN. The code used for the application of the single-cell transcriptomic ageing clock is available at https://github.com/SZTAKI-SU-Rejuvenation-Group/direct_reprogramming_dpsc_clock/blob/main/apply_inhibitory_neuron_clock.py.

## ACKNOWLEDGEMENTS

We are thankful to Johan Jakobsson, Janelle Drouin-Ouellet, Giorgia Tisoni, Yogita Sharma and all members of the Molecular Neurogenetics Laboratory at Lund University, and of the HCEMM-SU Neurobiology and Neurodegenerative Diseases Research Group. We thank Pálma Anna Zsolnai for her valuable assistance with lentiviral vector production at the Semmelweis Viral Vector Core. We are grateful to Nagy Nándor and Ádám Soós from the Laboratory of Stem Cell and Experimental Embryology (Nagy Lab), Department of Anatomy, Histology and Embryology, Semmelweis University, for providing the SSEA3 antibody. We thank the MultiPark FACS Core Facility and particularly Anna Hammarberg for the FACS analyses used in sorting single nuclei. We additionally acknowledge the Semmelweis FACS Core Facility for supporting all remaining FACS sorting procedures.

## AUTHOR CONTRIBUTIONS

K.K., G.V., A.F. and Á.Z. provided the human DPSCs used in the study. A.A.A., C.M. and K.P. designed and visualized the figures. C.M. led the formal analysis, data curation and computational framework for all bioinformatics work. A.A.A. and A.F. designed and performed the experimental work together with B.S., K.V., B.K., and Á.V. M.E.G., A.S. and K.L. designed, supervised and performed patch-clamp electrophysiological measurements. Z.D. performed the MS experiments and pre-processed the data, and C.M. performed the data analysis. K.V., J.G.J., A.A.A., Á.V. performed single nuclei sequencing experiments designed and supervised by C.M. and K.P. and analyzed by C.M. C.M and B.V. applied the epigenetic clocks under the supervision of C.K. and K.P. C.M. analysed the DNA methylation array data. E.Z.P. applied the single-cell transcriptomic clocks on the single-nuclei RNA sequencing data and performed the related analysis together with C.M under the supervision of C.K. and K.P. B.S., C.M. and A.A.A. wrote the manuscript with support from K.P. and with input from all authors. K.P. and A.F. conceived the study and oversaw overall direction and planning together with C.M. K.P. supervised the project. All authors provided critical feedback and helped shape the research, analysis and manuscript.

## ETHICS STATEMENT

All experiments involving human wisdom teeth samples and dental pulp stem cells described in this study were conducted under the ethical approval number (25459-4/2019/EKU). Human dermal fibroblasts used under ethical approvals REC 09/H0311/88 and IV/2625-1/2021/EKU.

## SUPPLEMENTARY INFORMATION

### Supplementary Data

The following data are provided in a separate .zip file:

**Supplementary Data 1:** Significantly differentially expressed protein-coding genes identified from bulk RNAseq data using DESeq2 and Benjamini-Hochberg correction, comparing NCAM1+ DPSC-iNs with DPSCs (n = 4).

**Supplementary Data 2:** Biological processes significantly enriched for protein-coding genes differentially expressed in bulk RNAseq data when comparing NCAM1+ DPSC-iNs with DPSCs (n = 4).

**Supplementary Data 3:** Pre-processed Mass Spectrometry-based proteomics data of DPSCs and DPSC-iNs (n = 4).

**Supplementary Data 4:** Significantly differentially abundant proteins identified from MS-based proteomics data using limma and Benjamini-Hochberg correction, comparing DPSC-iNs with DPSCs (n = 4).

**Supplementary Data 5:** Reactome pathways significantly enriched for proteins differentially abundant in MS-based proteomics data when comparing DPSC-iNs with DPSCs (n = 4).

**Supplementary Data 6:** Significantly differentially methylated protein-coding gene promoters identified from EPICv2 DNAm data using limma and Benjamini-Hochberg correction, comparing DPSC-iNs with DPSCs (n = 4).

**Supplementary Data 7:** DNA methylation clock predictions in DPSCs and DPSC-iNs (n = 4) and Pearson’s correlation test results.

**Supplementary Data 8:** Top cluster markers identified in snRNAseq data of day 17 DPSC-iNs (n = 3), using Wilcoxon Rank-sum test and Benjamini-Hochberg correction.

**Supplementary Data 9:** Significantly differentially expressed genes in lineage trajectories identified using tradeSeq in snRNAseq data of day 17 DPSC-iNs (n = 3).

**Supplementary Data 10:** Biological processes significantly enriched for genes differentially expressed in lineage trajectories in snRNAseq data of day 17 DPSC-iNs (n = 3).

**Supplementary Data 11:** Biological processes significantly enriched for Single-cell Inhibitory Neuron clock features (Muralidharan et al. 2025) having non-zero expression values in snRNAseq data of day 17 DPSC-iNs (n = 3).

**Supplementary Data 12:** Single-cell Inhibitory Neuron clock (Muralidharan et al. 2025) predictions in snRNAseq data of day 17 DPSC-iNs (n = 3).

## Supplementary Figures

Available in a separate .pdf document.

---

**Supplementary Figure 1.**
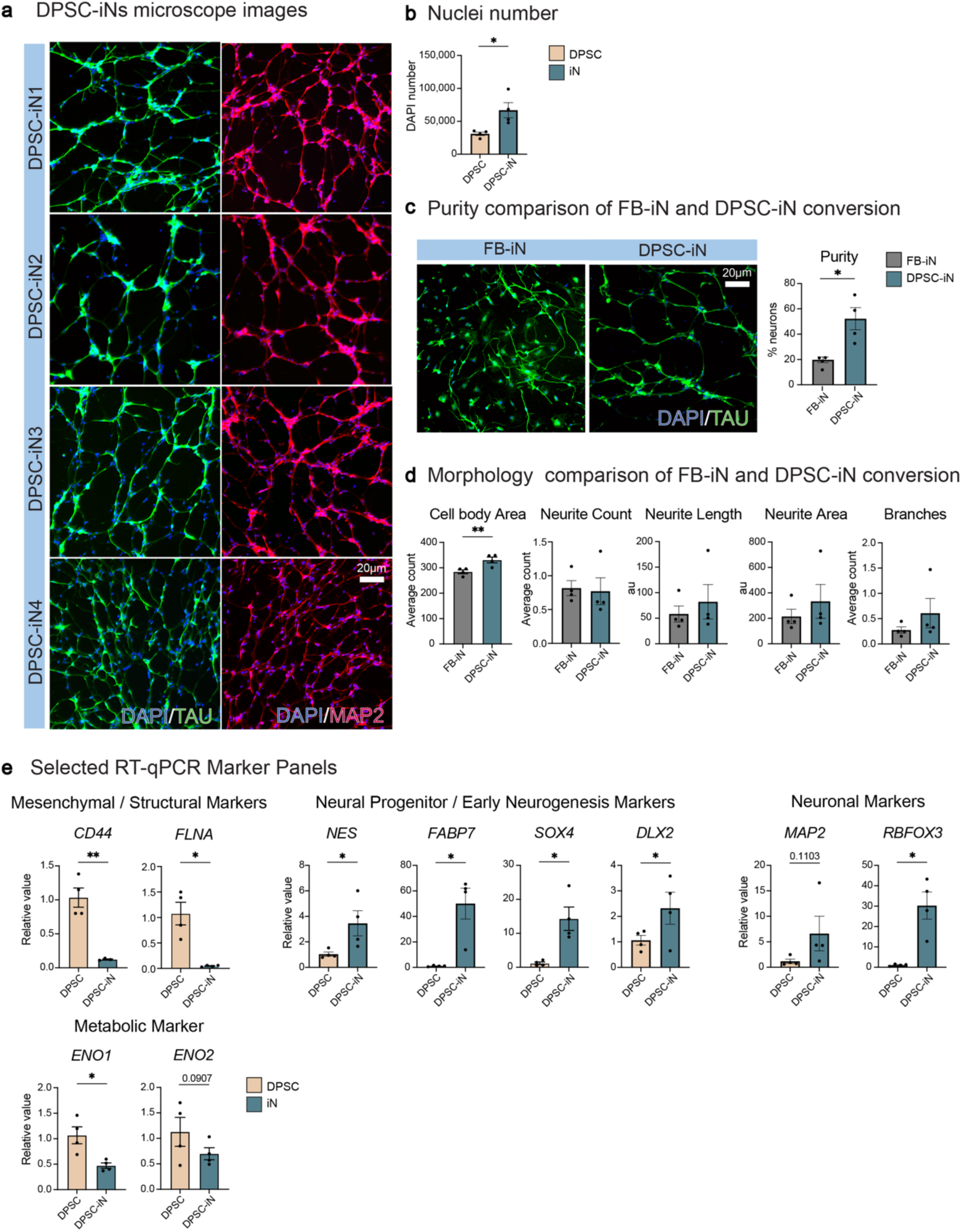
Characterization of DPSC-iN conversion across donors. Related to Figure 1**. a:** Representative ICC images from all four donor-derived DPSC-iN cultures showing consistent induction of neuronal markers (MAP2, TAU). **b:** Comparison of DAPI-positive nuclei numbers between parental DPSCs and donor-matched DPSC-iN cultures (n= 4 donors). **c:** Comparison of neuronal purity (% TAU⁺/DAPI⁺) between fibroblast-derived iNs and DPSC-derived iNs, showing significantly higher purity in DPSC-iNs (n= 4 donors for DPSC-iNs and n = 4 age-matched donors for FB-iNs). **d:** HCA-based morphological comparison of fibroblast-iNs and DPSC-iNs showing similar neurite number, neurite length, branch points and neurite area in DPSC-iNs (n= 4 donors for DPSC-iNs and n = 4 age-matched donors for FB-iNs). **e:** RT-qPCR analysis of different markers. DPSC-iNs showed increased expression of neural progenitor and neuronal genes (*NES, FABP7, SOX4, DLX2, MAP2,* and *RBFOX3*) and reduced expression of mesenchymal/structural and metabolic markers (*CD44, FLNA, ENO1,* and *ENO2*) compared with DPSCs (n = 4 donors). Scale bar: 20 μm. All results are shown as individual values with mean ± SEM from four independent donors (n = 4). Panels b and e were analyzed using paired two-tailed t-tests, whereas panels c and d were analyzed using unpaired two-tailed t-tests. Significance levels were defined as *p < 0.05, **p < 0.01, ***p < 0.001.

**Supplementary Figure 2.**
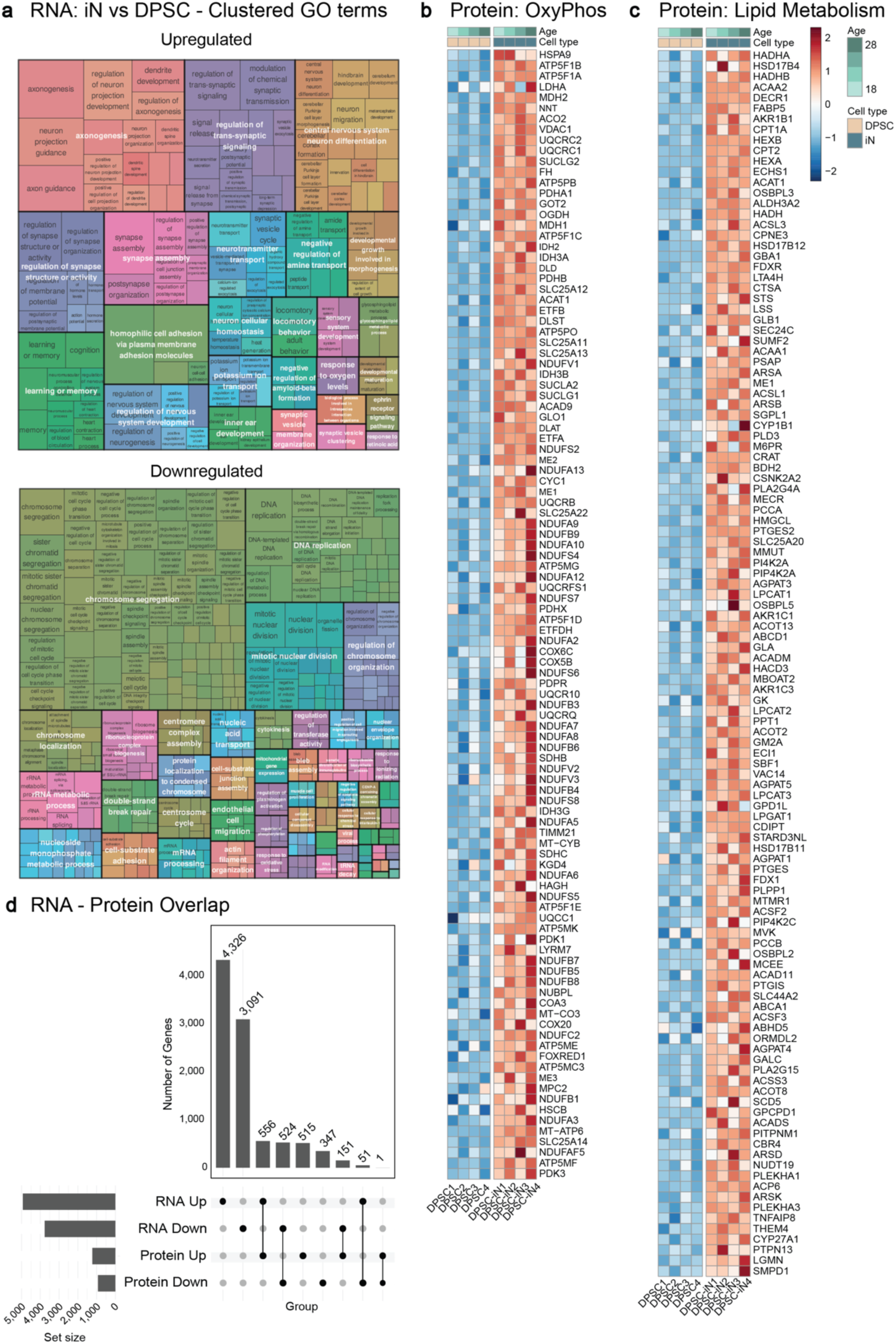
Biological processes supporting neuronal identity of DPSC-iNs. Related to Figure 2**. a:** Semantically grouped biological processes enriched for genes (left) upregulated and (right) downregulated in DPSC-iNs^#^ when compared to DPSCs in the bulk RNA-seq data (n = 4). **b:** Heatmap of proteins associated with oxidative phosphorylation showing consistent upregulation across all four DPSC-iN donors. **c:** Heatmap of lipid-metabolism-related proteins demonstrating increased abundance in DPSC-iNs compared with parental DPSCs. **d:** Plot showing number and overlap of features significantly upregulated or downregulated in the bulk RNA-seq and MS-based proteomic data of DPSC and DPSC-iN cells (n = 4), indicating major transcriptomic changes reflected at the proteomic level. Statistical significance was determined using DESeq2 (bulk RNA-seq data) or limma (MS-based proteomic data) coupled with Benjamini-Hochberg correction at false-discovery rate of 5%. ^#^ For bulk RNA-sequencing, DPSC-iNs were FACS sorted to only include NCAM1+ cells (Refer Methods). * Significance at adjusted p-value < 0.05.

**Supplementary Figure 3.**
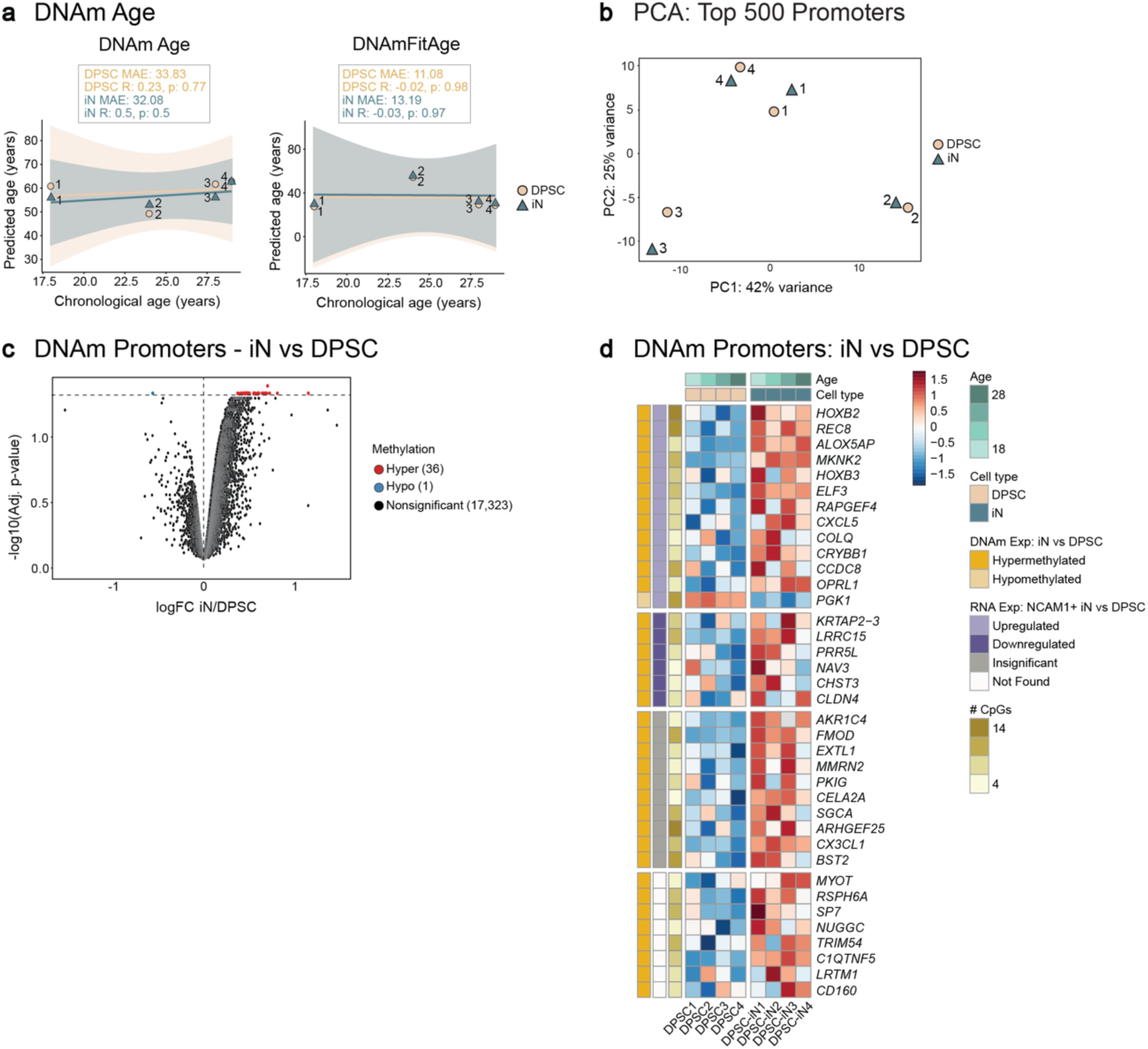
Alterations in promoter methylation during direct neuronal conversion. Related to Figure 2**. a:** Plot showing correlation of chronological age with predicted DNA methylation age based on (top) DNAm clock and (bottom) DNAmFitAge clock in the DNA methylation array data of DPSCs and DPSC-iNs (n = 4 donors). Using Pearson’s correlation test, the correlation coefficient R, and p-value were determined separately for the DPSCs and DPSC-iNs. **b:** Plots of principal components from PCA of top 500 (left) protein-coding gene promoter sites from DNAm data of DPSCs and DPSC-iNs across four donors (n = 4 donors). **c:** Volcano plot showing the distribution of log2 fold-change vs -log10 adjusted p-value of DNA methylation levels of gene promoter regions. Points highlighted in red were significantly hypermethylated, while those in blue were hypomethylated. Statistical significance was measured using limma coupled with Benjamini-Hochberg correction and false discovery rate of 5%. **d:** Heatmap showing promoter methylation changes between DPSCs and DPSC-iNs across donors. Significantly hypermethylated promoters are highlighted in yellow and the single hypomethylated promoter (*PGK1*) in blue.

**Supplementary Figure 4.**
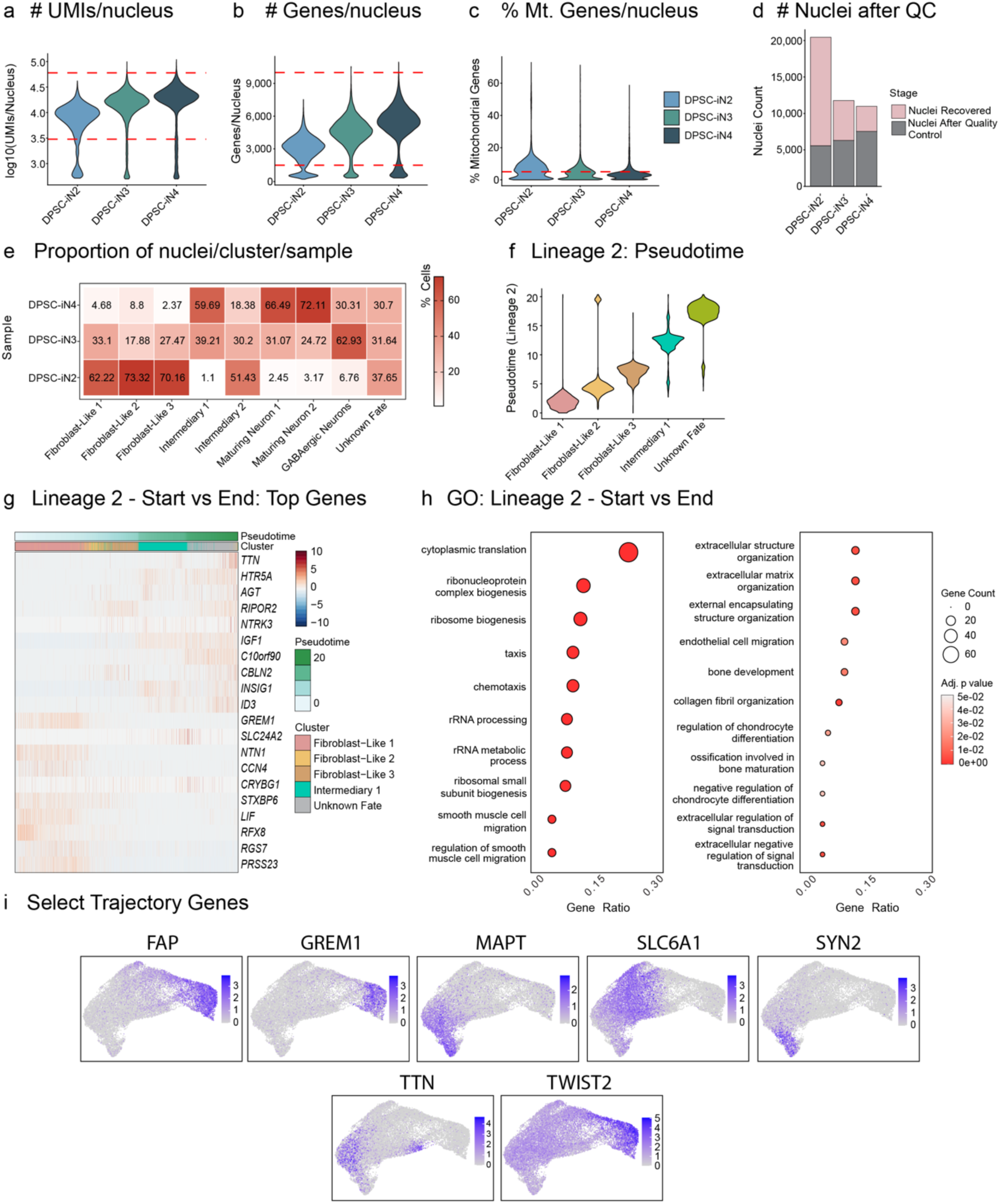
DPSC-iN snRNA-seq quality control and trajectory inference results. Related to Figure 4. a: Violin plots showing the number of UMIs per nucleus in the day 17 DPSC-iN snRNAseq data prior to quality control (n = 3). The dashed red lines indicate the filtering thresholds used during quality control. b: Violin plots showing the number of genes per nucleus in the day 17 DPSC-iN snRNAseq data prior to quality control (n = 3). The dashed red lines indicate the filtering thresholds used during quality control. c: Violin plots showing the percentage of mitochondrial genes expressed per nucleus in the day 17 DPSC-iN snRNAseq data prior to quality control (n = 3). The dashed red lines indicate the filtering thresholds used during quality control. d: Bar plot showing the proportion of nuclei recovered or detected by Cell Ranger, and that retained post quality control steps in the day 17 DPSC-iN snRNAseq (n = 3). e: Proportion of cells per cluster from each donor post quality control in the day 17 DPSC-iN snRNAseq data (n = 3). f: Violin plots showing the predicted pseudotime values per cluster across lineage 2 trajectory in the day 17 DPSC-iN snRNAseq. g: Heatmap showing the normalized and scaled expression of top 10 genes differentially expressed in the start or end points of the lineage 2 trajectory in the day 17 DPSC-iN snRNA-seq data (n = 3). h: Top biological processes enriched for genes differentially expressed between the start and end points of lineage 2 trajectory in the day 17 DPSC-iN snRNA-seq data (n = 3). i: UMAP plots showing the distribution of expression of selected genes, among the top differentially expressed genes across the lineage trajectories. Statistical significance was determined in g using tradeseq’s Wald test, and in h using ClusterProfiler, both coupled with Benjamini-Hochberg correction. False-discovery was controlled at corrected p-value < 0.05.

**Supplementary Figure 5.**
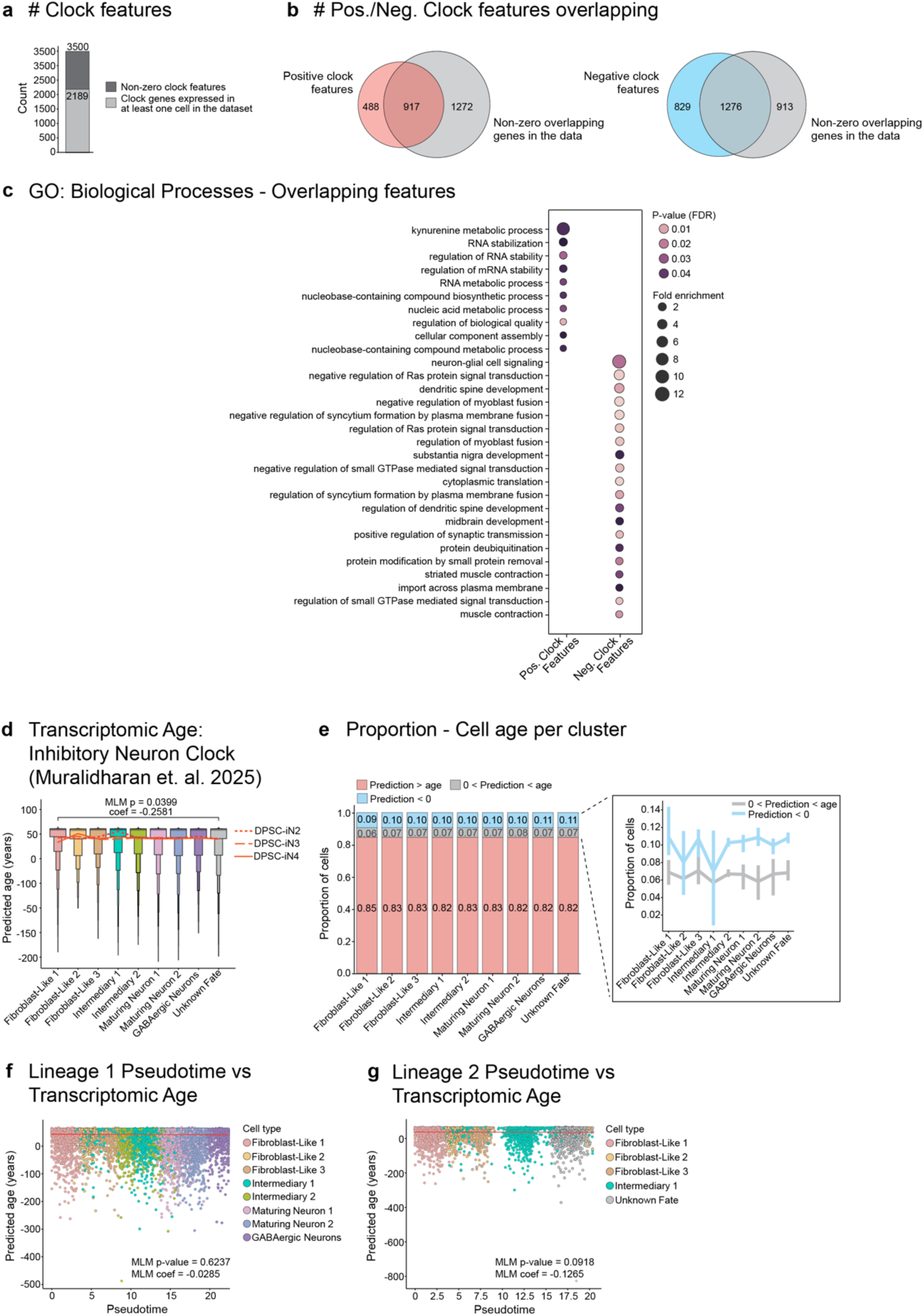
Single-cell Inhibitory Neuron clock predictions in DPSC-iNs. Related to Figure 4**. a:** Bar plot showing the proportion of Single-cell Inhibitory Neuron clock features with non-zero coefficient in at least one round of cross-validation^60^, and the proportion of those features expressed in the day 17 DPSC-iN snRNAseq dataset (n = 3). **b:** Plots showing the overlap of non-zero clock features expressed in the day 17 DPSC-iN snRNAseq dataset with the non-zero clock features having (left) an average positive coefficient and (right) a negative coefficient. **c:** Plot showing the top biological processes enriched for clock features with average positive or negative coefficients, and expressed in the day 17 DPSC-iN snRNAseq dataset. Significance was measured at FDR < 0.05. **d:** Plot showing the predictions of Single-cell Inhibitory Neuron clock in the day 17 DPSC-iN snRNAseq dataset (n = 3). Statistical significance was measured using a mixed linear model fit on the predicted age and chronological age with cluster and sex as additional variables followed by a pair-wise comparison between clusters. FDR < 0.05 was considered significant, and no significant results were found. **e:** Proportion of cells per cluster in the day 17 DPSC-iN snRNAseq dataset with predicted age above the chronological age, below but positive, or negative. **f and g:** Plots showing relationship, if any, between the predicted transcriptomic age and the predicted pseudotime of (f) lineage 1 trajectory and (g) lineage 2 trajectory in the day 17 DPSC-iN snRNAseq dataset. A mixed linear model was fit on to the predicted age and pseudotime of respective lineages, with cluster and sex as additional variables, and the donor as the grouping variable.

